# A disinhibitory basal forebrain to cortex projection supports sustained attention

**DOI:** 10.1101/2024.07.22.604711

**Authors:** Shu-Jing Li, Balazs Hangya, Unmukt Gupta, Kyle B. Fischer, James Fitzhugh Sturgill, Edward M. Callaway, Adam Kepecs

## Abstract

Sustained attention, as an essential cognitive faculty governing selective sensory processing, exhibits remarkable temporal fluctuations. However, the underlying neural circuits and computational mechanisms driving moment-to-moment attention fluctuations remain elusive. Here we demonstrate that cortex-projecting basal forebrain parvalbumin-expressing inhibitory neurons (BF-PV) mediate sustained attention in mice performing an attention task. BF-PV activity predicts the fluctuations of attentional performance metrics ― reaction time and accuracy ― trial-by-trial, and optogenetic activation of these neurons enhances performance. BF-PV neurons also respond to motivationally salient events, such as predictive cues, rewards, punishments, and surprises, which a computational model explains as representing motivational salience for allocating attention over time. Furthermore, we found that BF-PV neurons produce cortical disinhibition by inhibiting cortical PV+ inhibitory neurons, potentially underpinning the observed attentional gain modulation in the cortex. These findings reveal a disinhibitory BF-to-cortex projection that regulates cortical gain based on motivational salience, thereby promoting sustained attention.

**HIGHLIGHTS:**

- BF-PV activity predicts attentional performance metrics: reaction time and accuracy
- BF-PV responses reflect the computation of motivational salience-guided attention allocation
- Optogenetic activation of BF-PV neurons improves attentional performance
- BF-PV neurons produce cortical disinhibition through topographic projections and mediate gain modulation

## INTRODUCTION

The ability to sustain attention amidst competing sensory inputs is a cornerstone of cognitive faculty, crucial for survival and navigating life^1–3^. Attention is not a unitary construct but rather multi-faceted processes organized hierarchically. Consider a hunter in a forest, needing to maintain attention for hours tracking prey and evading predators. This attentional process begins with arousal (alerting), a state of heightened sensitivity to incoming stimuli. Next comes engagement (orienting), involving sustained focus on intricate tasks such as detecting subtle movements of prey or signs of danger. At the highest level, executive attentional control governs thoughts and actions according to internal goals. This includes focusing on specific locations where prey may hide (‘spatial attention’) and filtering for particular characteristics like a deer’s antlers through dense foliage (‘feature-based attention’)^4,5^. However, maintaining attention over time is mentally taxing, and lapses can expose the hunter to missed opportunities and potential threats. Such attentional lapses disrupt behavior in various disorders, ranging from Alzheimer’s and schizophrenia to attention-deficit/hyperactivity disorder (ADHD)^6–9^, underscoring the critical role of sustained attention in daily functioning and the necessity of understanding its neural underpinnings.

Studying attention presents a challenge because the term “attention” refers to both the cognitive processes involved and the resulting phenomenon, typically defined as enhanced selectivity in discerning signals amid noise^10^. This broad definition encompasses a diverse array of cognitive processes under the umbrella of “attention” and attempts to unify these under a single coherent theory have not succeeded. Additionally, research on attention has largely followed psychological typologies and assumed exhaustive sub-functions, but often underestimates commonalities of the underlying processes. Traditionally, alerting, orienting, and executive attention were considered largely distinct networks^11–13^. However, neuroimaging and neural recordings have revealed overlapping brain regions such as the frontal cortex, parietal cortex, norepinephrine (NE), and acetylcholine (ACh) neuromodulator systems^14–17^. This controversy underscores the need to understand these processes at a mechanistic level. In the hierarchy of attentional processes, arousal states can be studied through indicators like pupil dilation^18,19^, whereas at the high level, feature-based attention has been studied by examining how attending to specific sensory features modulates neural responses in relevant sensory areas^1,5,20,21^. Between arousal and feature-specific attention lies the allocation of attention over time, often simplified as heightened activity during good signal detection events, yet the precise computations remain elusive. Notably, attention exhibits remarkable temporal fluctuations, impacting performance across various sustained attention tasks^22–24^. Analyzing trial-to-trial variability offers insights into moment-to-moment attention dynamics, potentially helping to disambiguate attention from other cognitive variables.

Classic studies have implicated the basal forebrain (BF), with its extensive cortical projections, in attention, arousal, memory, and learning^25–29^, providing valuable insights into the neural basis of sustained attention. However, delineating its precise cognitive roles has been challenging. Lesion and pharmacological studies of the basal forebrain lacked cell-type specificity and temporal precision^27,30–32^, limiting their ability to isolate moment-to-moment performance fluctuations. The widely used five-choice serial reaction time task (5CSRTT) in rodent studies, while informative, does not effectively distinguish between sustained attention and related cognitive processes such as arousal and impulsivity^33,34^. The basal forebrain is primarily known for its cholinergic neural population, implicated in cognitive disorders like Alzheimer’s disease^35–37^. Recent recordings show that cholinergic signals primarily reflect arousal^29,38,39^ and reinforcement surprise^40–44^, do not track moment-to-moment attentional metrics^44^. Similarly, norepinephrine (NE) from the locus coeruleus is linked more to arousal/alertness and action execution than sustained focus^2,17^. Thus, while ACh and NE are crucial for arousal and alertness, they do not fully support sustained attention, suggesting the need to explore other neural substrates for this function.

This insight prompted us to investigate the other major output population in the BF, the parvalbumin-expressing GABAergic neurons (BF-PV), which also project to the cortex^45–47^ but have been largely overlooked in the context of sustained attention. Recent studies showed that optogenetic activation of BF-PV neurons promotes arousal^29,48^ and amplifies stimulus-driven cortical gamma oscillations^49–51^. While these findings implicate BF-PV neurons in regulating cortical states, their role in attention and other cognitive functions remains unknown.

In this study, we explore the role of BF-PV neurons in sustained attention by systematically examining their circuitry, behavioral functions, and computational features. We combine anatomical mapping, electrophysiological and optical recordings, and manipulation during an attention-demanding task, combined with computational modeling to interpret our results. Our findings reveal that BF-PV neurons project widely to cortical regions in a topographic manner and their activity closely tracks attentional performance dynamics. We demonstrate that these neurons compute ‘motivational salience,’ crucial for guiding sustained attention, and directly influence attentional performance. Furthermore, we identify a neural circuit mechanism whereby BF-PV neurons target cortical inhibitory neurons, leading to cortical disinhibition and sensory gain modulation. These results elucidate the neural circuitry and computations underlying sustained attention, providing new insights into the cell-type-specific contributions of the basal forebrain to cognitive function.

## RESULTS

### Topographic projections of BF-PV neurons to the cortex

The GABAergic parvalbumin-expressing (BF-PV) neurons are another major ascending system in the basal forebrain (BF) in addition to the region’s characteristic cholinergic neurons^45–47^, but it is unclear how their cortical projections are organized to support their cognitive functions. Therefore, we began by mapping their output connections to the cortex. We injected Cre recombinase-dependent adeno-associated virus (AAV) expressing fluorescent proteins into the BF of PV-Cre mice and identified axonal fibers in the cortex. We observed that PV neurons at the anterior end of the BF (aBF-PV) predominantly project to midline cortical structures such as the medial prefrontal cortex (mPFC) and retrosplenial cortex (RSC) **(Figure 1A-C)**. In contrast, PV neurons at the posterior end of the BF (pBF-PV) preferentially project to more lateral cortical regions including the motor cortex (M1/M2), somatosensory cortex (SS), and auditory cortex (ACx) **(Figure 1D-F)**. This separation of cortical targets between anterior and posterior BF-PV pools may facilitate finer control of the cortex during cognitive functions, in contrast to uniformly broadcasting via diffuse projections.

**Figure 1.**
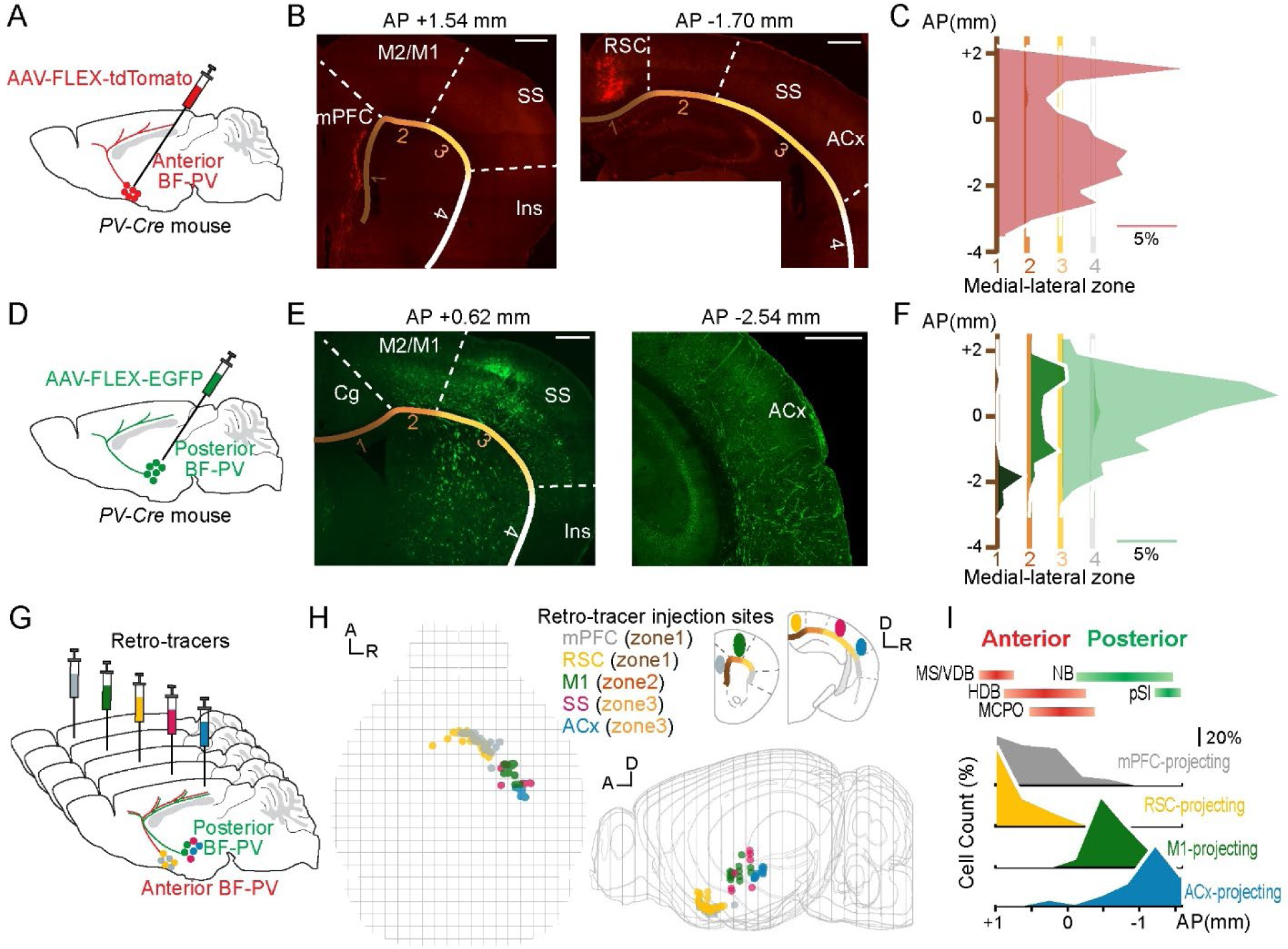
Cortically-projecting basal forebrain parvalbumin-expressing inhibitory neurons (BF-PV) show topographic organization. (A-F) Mapping the projections from anterior (A-C) and posterior (D-F) BF-PV neuronal pools to the cortex. (A & D) Schematic of anterograde viral tracing approach. (B & E) Labeled projections in various cortical regions. The labeled axon terminals are counted in each cortical zone divided by medial-lateral positions (zones: 1, medial; 2 & 3, dorsal-lateral; 4, ventral-lateral). mPFC, medial prefrontal cortex; M2/M1, secondary and primary motor cortex; SS, somatosensory cortex; Ins, insular cortex; RSC, retrosplenial granular cortex; Cg, cingulate cortex; ACx, auditory cortex. Scale bar, 500μm. (C & F) Distribution of the cortical projections from anterior (C) and posterior (F) BF-PV pools are separated, which is enriched in medial (zone 1) and lateral cortex (zone 2 & 3), respectively. Data is presented as the percentage of labeled axon terminals located in each medial-lateral zone and anterior-posterior coronal plane in individual mice and averaged across animals (n = 2 in (C), n = 3 in (F)). Scale bar, 5%. (G-I) Mapping the locations of cortical-projecting PV neurons in different BF sub-regions. Schematic of retrograde viral tracing approach. 3D view of the locations of BF-PV neurons that target different cortical regions. A, anterior; D, dorsal; R, right. The distribution of BF-PV neurons is organized based on cortical targets, with anterior BF-PV targeting medial cortices whereas posterior BF-PV targeting dorsal-lateral cortical areas. Data is presented as the percentage of labeled PV neurons located in each anterior-posterior coronal plane in individual mice and averaged across animals (n = 4 for tracing from mPFC, RS, and ACx individually; n = 3 for tracing from M1). Red and green bars on top represent the anterior-posterior distance each sub-region covers. MS, medial septum nucleus; VDB, vertical diagonal band; HDB, horizontal diagonal band; MCPO, magnocellular preoptic nucleus; NB, nucleus basalis; SI, substantia innominata. Scale bar, 20%. See also Figure S1.

The sub-regions of the basal forebrain have previously been loosely delineated based on anatomical landmarks and topography of cholinergic cell groups with different projection targets^26,52^, but whether BF-PV neurons also follow the same spatial organization is unknown. Therefore, we investigated the organization of BF-PV neurons in different basal forebrain sub-regions with respect to their cortical projection targets. We injected retrograde tracers into cortical areas that differ in medial-lateral positions, then identified the cell types of labeled neurons in the basal forebrain and mapped their locations onto an annotated 3D spatial framework based on the Allen Brain Atlas^53^ **(Figure 1G-H, Figure S1)**. Consistent with the anterograde tracing results, BF-PV neurons projecting to the medial cortical regions like the prefrontal cortex (mPFC) and retrosplenial granular cortex (RSC) were mainly found in the anterior BF sub-regions covering the medial septum nucleus (MS), vertical and horizontal diagonal band (VDB, HDB), and the magnocellular preoptic nucleus (MCPO), with their density decreased from rostral to caudal **(Figure 1H-I, Figure S1A-C)**; while those neurons projecting to lateral cortical regions like the motor cortex (M2/M1), somatosensory cortex (SS), and auditory cortex (ACx) were primarily found in the posterior BF sub-regions spanning the nucleus basalis (NB) and the substantia innominata (SI) **(Figure 1H-I, Figure S1D-G)**. In addition, although BF-PV and cholinergic neurons were both found in these sub-regions, their locations and densities within individual sub-regions were shifted. BF-PV neurons are located more medially in MS but more laterally in HDB than cholinergic neurons **(Figure S1B)**. Some cortical regions, like mPFC, received enriched inputs from BF-PV neurons more than from cholinergic neurons **(Figure S1A-C)**. These findings indicate that an accurate delineation of BF sub-regions would require evaluation of both neuron types.

Together, these results show that BF-PV neurons send extensive projections to the cortex, albeit with a slightly different sub-regional distribution compared to cholinergic neurons^54^. Their cortical projections follow a topographic organization: those in the anterior BF sub-regions primarily target medial cortical areas, while those in the posterior BF sub-regions focus on more lateral cortices. This suggests a structured hierarchy of BF-PV cortical projections, similar to the cholinergic system^26,55,56^, which provides a potential neural substrate for both rapid signal broadcasting cortex-wide and region-specific regulation to support attentional processes.

### BF-PV neural spiking activity predicts behavioral correlates of sustained attention

The functions of the basal forebrain have been implied in attention and learning through its vast projections to the cortex^26–29^. Although GABAergic PV neurons constitute a major cortically-projecting BF population, their specific cognitive roles remain unknown. To explore this question, we employed an auditory detection task for head-fixed mice that involves sustained attention and reinforcement learning, and allows temporal separation of these behavioral variables^44^ **(Figure 2A)**. In this task, mice had to detect and respond to a specific “target” tone that was masked by a noisy background stream. Each trial began with a start signal. Then during the subsequent task epoch, called the *foreperiod*, mice were required to refrain from licking the waterspout until the delivery of “target” or “non-target” tones (stimulus). Successful rapid lick response to the “target” tone was considered a Hit and resulted in a water reward (outcome), while licking to the “non-target” tone was considered a False alarm and led to air-puff punishment. To increase the attentional load and to discourage impulsive licking behaviors, the tones were delivered at unexpected moments (foreperiod duration was randomized within 0.1-5s, exponential distribution), and animals’ premature licks before tone onset canceled the trial. The stimulus detection difficulty was graded by varying the loudness of the tones relative to the background noise (SPL, relative sound pressure level randomized within 10-50 decibels). Thus this task requires animals to keep attentional effort during the foreperiod in preparation to perform rapid licks upon “target” tones that are difficult to predict or detect **(Figure S2A)**. After training, mice performed the task well: their behavioral accuracy (Hit rate) and discriminability score in detecting the “target” tone increased **(Figure 2B)**, while reaction time (RT) decreased **(Figure S2B)** as stimulus detection difficulty reduced (SPL increased).

**Figure 2.**
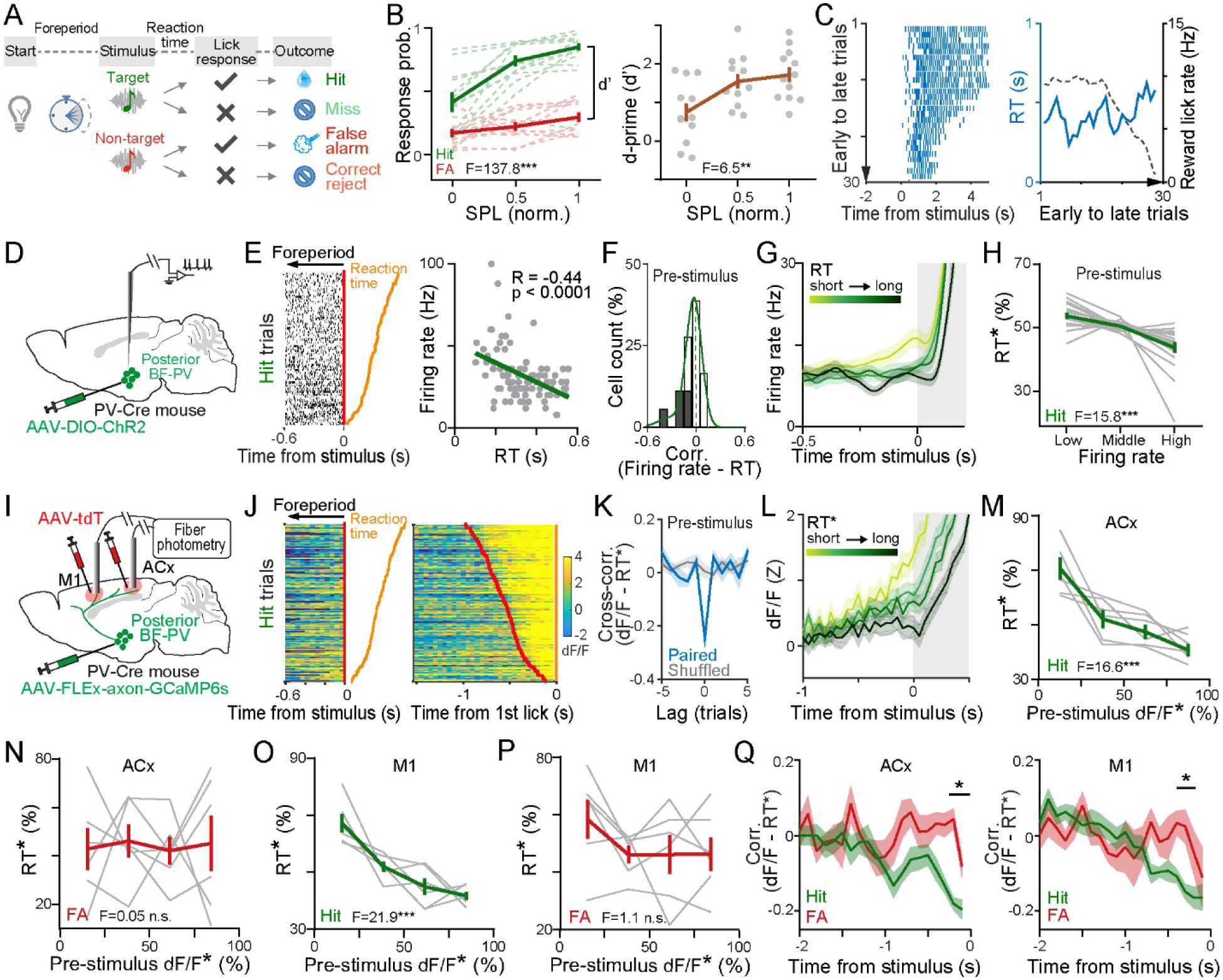
BF-PV activity predicts reaction time in sustained attention task. (A-C) A head-fixed auditory detection task to measure sustained attention performance. Trial structure and possible outcomes. Mice were required to withhold licking after the start signal during a variable foreperiod, their lick responses to “target” or “non-target” stimuli (pure tones of well-separated pitch but varying intensity in white noise) resulted in water reward (Hit) or air-puff punishment (False alarm, FA), respectively. No feedback was given for trials with no or slow (> 1s) licking (Miss or Correct reject). Psychometric curves (left) and detection discriminability (right) as a function of detection difficulty (SPL, normalized sound pressure level; 1, loudest; 0.5, intermediate; 0, weakest) for individual mice (light curves or gray dots) and grand average (thick curves, n = 12 mice). Left, lick probability to “target” versus “non-target” stimulus [Hit vs FA, F(1, 11) = 137.8, p < 0.0001; In Hit trials, SPL 0 vs 0.5, p = 0.0006, SPL 0.5 vs 1, p = 0.018; In FA trials, SPL 0 vs 0.5 vs 1, n.s.]. Right, the d-prime score (d’) shows the distance between the Hit and FA rate [F(2, 33) = 6.5, p = 0.004]. (A) Fluctuations of reaction time (RT) across trials. Left, raster plot of licks in an example session. Right, traces of RT and lick rate to water reward (sliding window, 5 trials) from early to late in a session (Hit trials at same SPL are presented). (D-H) Spiking activity of BF-PV neurons. (B) Approach schematic for electrophysiological recordings of neural spikes from identified PV neurons located in posterior BF (n = 18 neurons collected from 4 mice). (C) Reaction time-predicting single neuron. Left, spike raster during the foreperiod before stimulus onset (red line) for Hit trials sorted by reaction time (orange ticks). Right, pre-stimulus firing rate (in a 250ms window preceding stimulus onset) negatively correlates with the following reaction time [R = -0.44, p < 0.0001, Pearson]. Each dot corresponds to one trial. (D) Distribution of correlation coefficient between pre-stimulus firing rate and reaction time (as R in (E)) for individual neurons. The shaded represent neurons with significant correlation [p < 0.05, Pearson]. A non-linear fitting (Gaussian, green) shows asymmetrical distribution with a tail to the left [Skewness= -1.8]. (E) Peri-event time histograms (PETHs) aligned to stimulus onset for Hit trials from an example neuron grouped by reaction time duration. Spiking activity is increased prior to stimulus onset. (F) Average of reaction times (rank, RT*) in Hit trials grouped by low, middle versus high pre-stimulus firing rate (split in each session) for individual neurons and grand average [F(2, 34) = 15.8, p = 0.0001, n = 18]. RT* was calculated as the percentile rank of reaction times in each session. Population spiking activity predicts reaction time. (I-Q) BF-PV axonal activity in the auditory (ACx) and motor cortex (M1). (G) Approach schematic for fiber photometry recordings in the cortex for axonal calcium activities of posterior BF-PV neurons (n= 6 mice, each animal contributes at least three sessions). (H) Example photometry raster before stimulus onset (left, red ticks) or first anticipatory lick (right, orange ticks) for Hit trials sorted by reaction time. Raising axonal activity during the foreperiod precedes the onset of stimuli or licks and predicts fast reaction time. (I) Cross-correlation analysis shows a negative correlation between pre-stimulus axonal activity (in a 500ms window preceding stimulus onset) and reaction time (Z-scored in each session) in the same trial. Gray line, shuffled trials; Shaded area, ± SEM across animals (n = 6). (J) Average cortical axonal activity aligned to stimulus onset for Hit trials from an example animal grouped by reaction time (rank). Axonal activity ramps up before stimulus onset. (M-P) Pre-stimulus axonal activity predicts reaction time. Average reaction times (rank) are tuned by pre-stimulus axonal activity levels (rank) in Hit trials [(M), ACx, F(3. 20) = 16.6, p < 0.0001; (O), M1, F(3, 20) = 21.9, p < 0.0001], but not in False alarm trials [(N), ACx, F(3, 20) = 0.05, p = 0.99; (P), M1, F(3, 20) = 1.1, p = 0.36] for individual mice (gray curves) and pooled across animals (green curve, n = 6). (Q) Average correlation coefficients between reaction time (rank) and axonal activities across time aligned to stimulus onset. Note that the negative correlation is more significant for Hit than False alarm trials prior to stimulus onset [100 ms time bins; ACx, Interaction between trial-types and time, F(19, 190) = 2.4, p = 0.0014; Hit vs FA for window (-0.3s -0.2s), p = 0.03; for window (-0.2s -0.1s), p = 0.0003; M1, Interaction, F(19, 190) = 3.2, p < 0.0001; Hit vs FA for window (-0.3s - 0.2s), p = 0.035]. Data are represented as means ± SEM. *p < 0.05, **p < 0.01, ***p < 0.001, n.s., not significant; using repeated measures one-way ANOVA for groups of three or more samples followed by Tukey’s multiple comparisons, and repeated measures two-way ANOVA for groups on two independent variables followed by Sidak’s multiple comparisons. See also Figure S2 & S7.

In humans, sustained attention can wander from moment to moment, and the momentary levels of attention can be read out by performance metrics such as reaction time and accuracy trial-by-trial^57–60^. Similarly, in mice performing a sustained attention task, a correlation between temporal attention and reaction time by varying expectancy has been found^44^, suggesting the same behavioral approach for analyzing momentary attention is applicable across species. Here, we also observed remarkable fluctuations in the reaction times across trials in mice doing this sustained attention-demanding auditory detection task **(Figure 2C)**, with a dynamic independent of reward-collecting behaviors driven by the motivation to consume water **(Figure S2C)**. Therefore, we hypothesized if BF-PV neurons track the fluctuations of attention, their activity dynamics would reflect a correlation with these attentional performance metrics. In addition, since in this task the sustained attention is allocated during the foreperiod for the detection of upcoming stimuli, we could operationalize attentional modulation as neural activity during the foreperiod that predicts reaction time (negative correlation) and/or accuracy (positive correlation). This helps to isolate the attention-related activity that precedes perceptual discrimination (sensory stimulus) and action execution (licking movement).

After establishing the measurement of sustained attention in this auditory detection task, we set out to investigate the relationship between sustained attention and BF-PV activity in behaving mice. We mainly focused on the auditory and motor cortex-projecting posterior BF-PV neurons, since these two cortical areas are critical for the auditory detection performance. We rendered these neurons light-sensitive by viral delivery of Cre-dependent channelrhodopsin (ChR2) in PV-Cre mice, and performed electrophysiological recordings of well-isolated single units **(Figure 2D)**. PV-expressing neurons were identified by optogenetic tagging approach^44,61,62^ based on their significant short latency action potentials to brief (1 ms) blue light pulses (n = 18 out of 154 units with restricted locations within the BF from 4 mice; p < 0.01; SALT test for optogenetic identification). **Figure 2E** shows the Hit trials from an example neuron with increasing firing rate during the foreperiod that was strongly correlated with short reaction time on a trial-by-trial basis. A substantial fraction of individual BF-PV neurons exhibited pre-stimulus activity that was negatively correlated with reaction time **(Figure 2F**, p < 0.05 in ∼28% of neurons). When viewed as a population, they displayed a remarkable firing rate increase during the foreperiod **(Figure 2G)**, with high activity predicting short reaction time **(Figure 2H, Figure S2E)**.

These results demonstrate the neural spiking activity of the BF-PV population predicts reaction time in the auditory detection task, which is exactly as we would expect to see in neurons that track attention fluctuations. Considering the anatomical findings that BF-PV neurons project to the cortex, these evidence indicate that BF-PV neurons are in a good position to mediate attentional modulation for auditory sensory detection, probably through their cortical projections. If that’s the case, the attention-related signal is expected to be transmitted through their axons to the relevant cortex.

### Axonal activity of BF-PV neurons in cortex shows correlates of sustained attention

To investigate whether the spiking activity we observed at BF-PV cell somata is conveyed through their projections to the cortex, and to understand what information the cortex receives, we directly measured BF-PV axonal activity in both auditory and motor cortex. We expressed a Cre-dependent axon-enriched calcium indicator axon-GCaMP6s^63^ in the posterior BF-PV neurons in PV-Cre mice through viral delivery and monitored axonal calcium transient in both cortices via fiber photometry **(Figure 2I, Figure S2F-G)**. Similar to the spiking data we observed, the axonal activity of BF-PV neurons also ramped up during the foreperiod, followed by a remarkable increase that was better aligned to stimulus onset than to licking onset **(Figure 2J).** This pre-stimulus axonal activity was inversely correlated to reaction time **(Figure 2K-L, Figure S2H-I)**, reliably predicting it **(Figure 2M, O)**, and this observation was consistent in both auditory and motor cortex across animals. Moreover, this negative correlation was time-specific, more pronounced near stimulus onset than earlier **(Figure 2Q)**, which is aligned with human and primate studies that temporal focus of attention accelerates the reaction times in the detection of upcoming stimulus^57,58,64^.

We then considered whether this faster reaction time-predictive activity could be caused by other plausible mechanisms such as higher motor vigor. The attentional modulation is supposed to promote the detection performance specifically for the “target” tone, such as reducing reaction time and increasing detection discriminability in Hit trials, whereas the motor vigor modulation would affect reaction time and lick response for both “target” (Hit) and “non-target” (False alarm) tones. The analysis of reaction times showed that the negative correlation between reaction time and BF-PV pre-stimulus activity was more evident in Hit compared to False alarm trials, suggesting it can’t be merely explained as a motor vigor correlate **(Figure 2M-Q)**. In addition, the analysis of trial outcomes showed a higher BF-PV activity during the foreperiod preceding correct tone detection (Hit) compared to error trials (False alarm or Miss; **Figure 3A-B**). Moreover, higher foreperiod activity was associated with increased accuracy on the psychometric curve **(Figure 3C)** and an upward shift on the discriminability curve **(Figure 3D)**. These results reveal that pre-stimulus BF-PV activity predicts reaction time and detection accuracy specifically for the “target” tone, aligning with selective attentional modulation rather than general vigor modulation.

**Figure 3.**
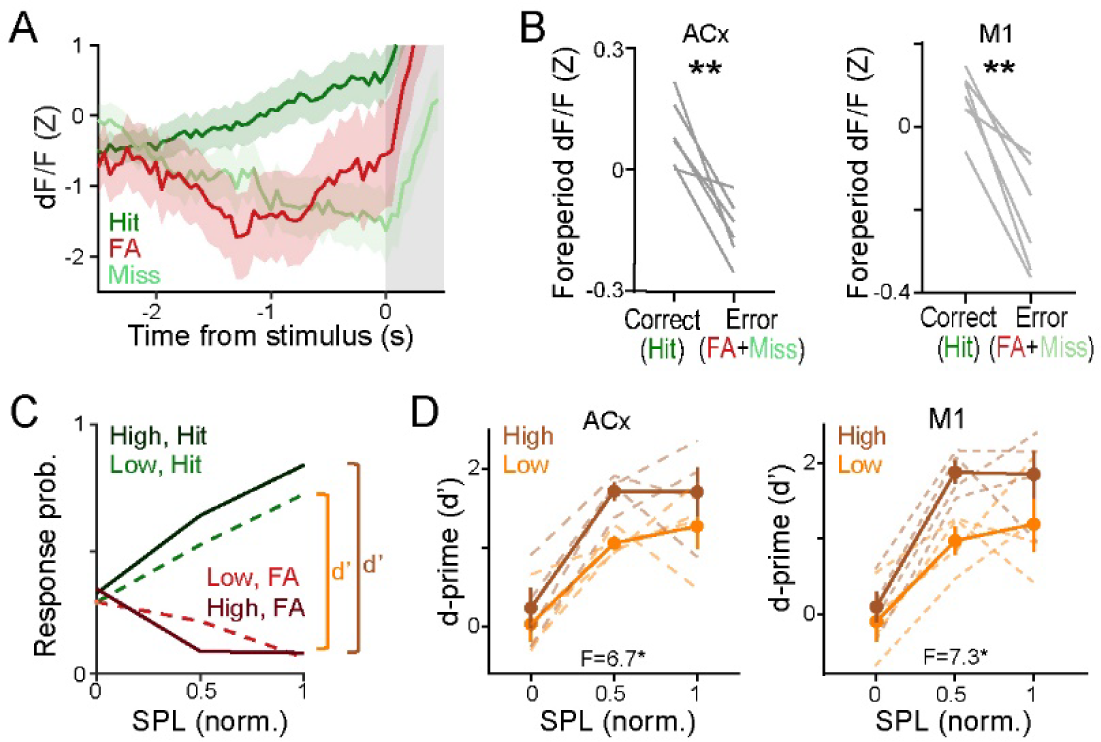
BF-PV activity predicts accuracy of sustained attention performance. (A) BF-PV axonal activity during the foreperiod for Hit, False alarm, and Miss trials aligned to stimulus onset in an example session. (B) Average axonal activity during the foreperiod is higher in correct (Hit) than error trials (False alarm + Miss) across animals [ACx, p = 0.004; M1, p = 0.002; n = 6; paired t-tests]. (C) Psychometric curves as a function of detection difficulty (SPL, normalized sound pressure level) are separated by High-versus Low-foreperiod activity (median-split) in an example animal. (D) Elevated foreperiod activity predicts higher detection discriminability. The d-prime score as a function of detection difficulty (SPL) is separated by High-versus Low-foreperiod activity for individual mice (light curves) and pooled across animals (thick curves) [ACx, High vs Low, F(1, 6) = 6.7, p = 0.04; High vs Low at SPL 0.5, p = 0.014; M1, High vs Low, F(1, 6) = 7.3, p = 0.035; High vs Low at SPL 0.5, p = 0.029; n = 4, mice with all SPL levels applicable for this analysis were included; repeated measures two-way ANOVA followed by Sidak’s multiple comparisons]. Data are represented as means ± SEM. *p < 0.05, **p < 0.01, ***p < 0.001, n.s., not significant. See also Figure S3 & S7.

We also considered the possibility that errors may be due to lower arousal since arousal influences attentive behaviors^17^. Therefore, we monitored arousal status by tracking pupil size as a proxy^65^ **(Figure S3A)**. We found pupil size increased slowly after both “target” and “non-target” stimuli regardless of outcomes **(Figure S3B-C)**, and pupil size before stimulus didn’t predict reaction time or discriminability **(Figure S3D-E)**. This is consistent with previous findings that arousal doesn’t solely dictate attention in a linear manner^17^. Notably, we found larger pupils linked to higher BF-PV activity during the baseline (4s-long window before stimulus onset) but not during the stimulus or outcome epochs **(Figure S3F)**, suggesting that arousal level may be associated with slow fluctuation of BF-PV baseline activity (at a time scale of seconds)^29,48^, but cannot explain the fast response properties of BF-PV neurons under behavioral task structure.

Together, these findings demonstrate that cortex-projecting BF-PV neurons track momentary attention levels, as elevated BF-PV activity is predictive of future reaction time and accuracy ― two behavioral correlates of sustained attention ― on a trial-by-trial basis.

### BF-PV responses track salient behavioral events

Psychological studies suggest that the allocation of attention can be influenced by multiple factors, such as motivation, value, salience, and surprise^22,66–68^. This led us to ask whether these phenomena can be reflected by BF-PV neural activity. To explore this, we used a battery of behavioral tasks to systematically study the activity pattern of BF-PV neurons in relation to stimulus presentation, outcome valence, and outcome surprise.

First, we examined the stimulus- and outcome-related responses of BF-PV neurons in the auditory detection task **(Figure 4A-G)**. The neural spikes of identified BF-PV neurons showed long latency and sustained activation after the delivery of reward and punishment **(Figure 4B-C)**, which contrasts with the fast and phasic cholinergic responses **(Figure S4A-B)**. Similarly, the axonal activities of BF-PV neurons in the auditory and motor cortex were also induced by both reward and punishment, as well as predictive cues **(Figure 4D-F, Figure S4C-D**). To test whether BF-PV neural responses to the sound stimuli are innate or acquired through learning, we looked at the changes in BF-PV activity at different training stages. The response to a sound stimulus was absent initially during the early learning stage, but acquired after mice learned the sound-reward association and developed anticipatory licking to the sound **(Figure S4E)**. This finding suggests that BF-PV’s neural response is developed through learning toward task-relevant stimuli that predict crucial outcomes. Furthermore, in the auditory detection task, the graded signal-to-noise ratio created varied stimulus detection uncertainty as reflected by graded Hit rates **(Figure 2B)**, thus resulting in different levels of outcome expectation and surprise. We observed that BF-PV axonal responses to punishment were not systematically modulated by stimulus strength, suggesting that false detections arise independent of the stimulus **(Figure S4F)**. In contrast, their responses to water reward increased, while the responses to stimulus decreased as the signal-to-noise ratio of stimulus dropped **(Figure 4G, Figure S4G)**. This inversely correlated pattern matches with how the levels of expectation and surprise are oppositely modulated by varied stimulus detection difficulty.

**Figure 4.**
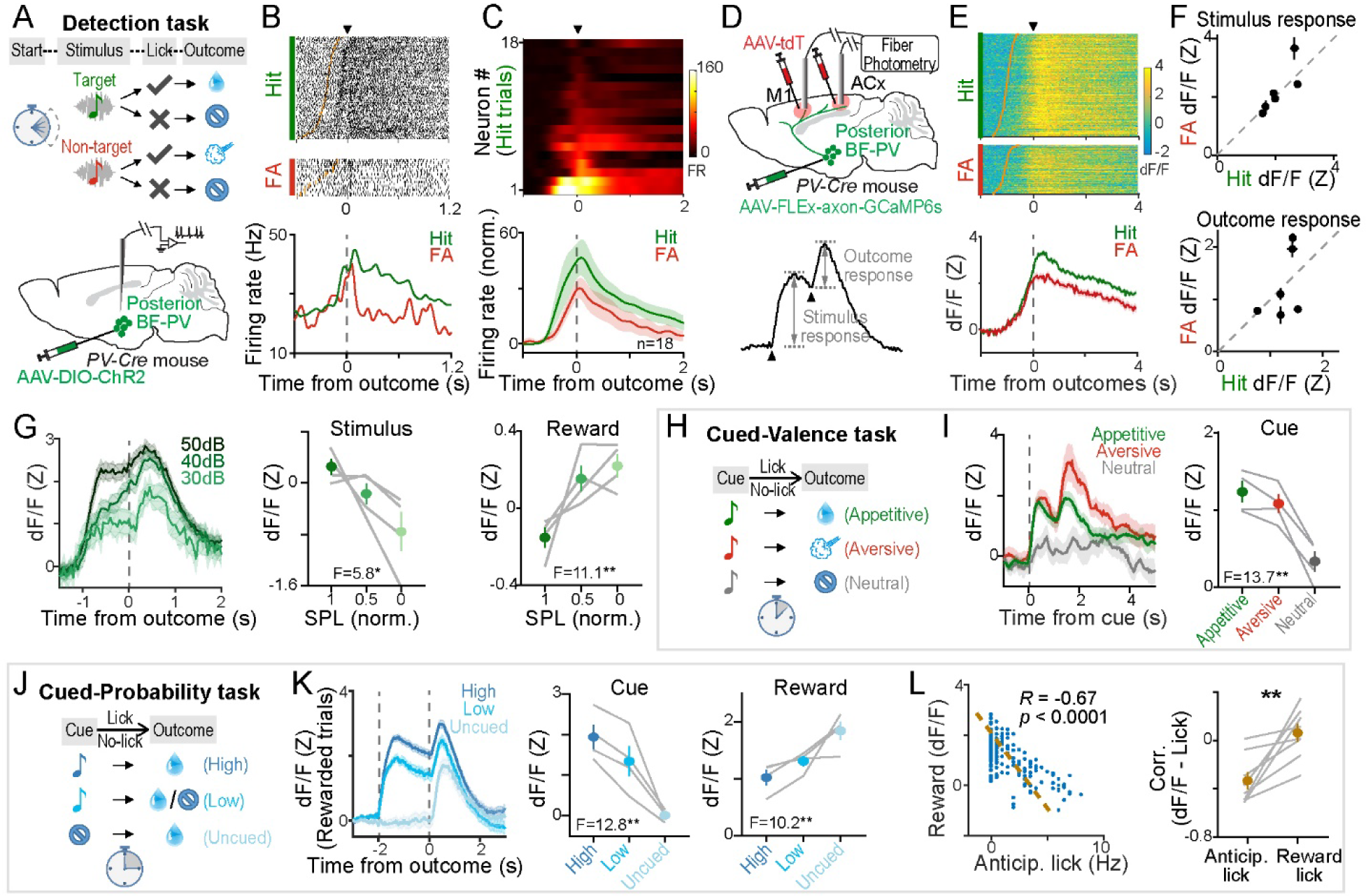
BF-PV neurons respond to significant behavioral events: predictive cues, outcomes, and surprises. (A-C) Spiking activity of BF-PV neurons in mice performing auditory detection task. (A) Schematics of trial structure (top) and approach for the electrophysiological recording of identified posterior BF-PV neurons (bottom, n = 18 neurons collected from 4 mice). (B) Spike raster (top) and peri-event time histograms (PETHs, bottom) aligned to outcome onset (arrowhead or dash line) and sorted by reaction time (orange ticks: stimulus onset). A single neuron shows positive responses to both water reward and air-puff punishment. (C) PETHs of individual neurons sorted by response amplitude (top) and average population activity (bottom). (D-G) BF-PV axonal activity in the auditory cortex in mice performing the auditory detection task. (D) Top, approach schematic for fiber photometry recordings in the cortex for the axonal calcium activities of posterior BF-PV neurons (n = 6 mice). Bottom, the magnitudes of stimulus or outcome response are measured as the changes before and after event onset (arrowhead). (E-F) Example axonal activity time series (E) and response magnitudes in individual mice (F) show positive responses to both reward and punishment outcomes [p = 0.996], as well as to both predictive stimuli [p = 0.598]. Photometry raster (E, top) and average plot (E, bottom) for Hit and False alarm trials are aligned to outcome onset and sorted by reaction time (orange ticks: stimulus onset). (G) Example axonal activity time series (left) and response magnitudes across animals (middle and right) show decreasing stimulus response [F(2, 9) = 5.8, p = 0.02] and increasing reward response [F(2, 9) = 11.1, p = 0.004] are tuned by reducing detection difficulty (normalized sound pressure level). Response magnitude is normalized to mean stimulus or reward responses within subject and shown in individual mice (gray curves) and grand average (green). n = 4, mice with all SPL levels applicable for this analysis were included. (H-L) BF-PV axonal activity in the auditory cortex in mice performing Cued-Valence task (H-I) and Cued-Probability task (J-L) (n = 4 mice). (H, J) Schematics for the trial structure. (I) Example axonal activity time series aligned to cue onset (left) and response magnitudes across animals (right) show positive responses to both reward- and punishment-predictive cues, but no response to cue predicting neutral outcome [F(2, 9) = 13.7, p = 0.0018]. (K) Example axonal activity time series (left) and response magnitudes across animals (middle and right) show decreasing cue response [F(2, 9) = 12.8, p = 0.002] and increasing reward response [F(2, 9) = 10.2, p = 0.005] are tuned by decreasing reward expectation (increasing surprise). The vertical dash lines represent the onsets of cue and reward. (L) Left, reward responses negatively correlate with the preceding anticipatory lick rate during cue epoch in an example session [R = -0.67, p < 0.0001, Pearson, each dot corresponds to one trial]. Right, reward responses show a greater negative correlation with anticipatory lick rate than with reward lick rate across animals [p = 0.006], reflective of reward expectation rather than movement. In each animal, conditions under high- and low-reward probability are separately evaluated. Data are represented as means ± SEM. *p < 0.05, **p < 0.01, ***p < 0.001, n.s., not significant; using paired t-tests for two groups and ordinary one-way ANOVA followed by Tukey’s multiple comparisons for groups of three or more samples. See also Figure S4 & S7.

To further quantitatively evaluate how the stimulus- and outcome-related responses of BF-PV neurons are scaled by varying value and surprise, we monitored their axonal activity in two additional Pavlovian tasks. In the “Cued-Valence” task, three distinct sound cues (different pitches, same loudness) precisely predicted either an appetitive (water reward), aversive (air-puff punishment), or neutral (null) outcome, thus carrying different levels of value **(Figure 4G)**. We observed positive responses to both appetitive and aversive cues, but not by the neutral cue **(Figure 4H-I, Figure S4H)**, indicating the stimulus and outcome responses of BF-PV neurons are valence-free and linked to the “salience/significance” related to the task. In the “Cued-Probability” task, two unambiguous sound cues (different pitches, same loudness) indicated a high (100%) or low (50%) probability of receiving a water reward; Additionally, water rewards were delivered without any preceding cue (uncued) in random trials, thereby creating varying levels of reward expectation and surprise **(Figure 4J)**. We observed larger responses for cues that predicted higher reward probability, and a reward response that was inversely proportional to reward expectation, with the largest response to the most surprising, uncued reward **(Figure 4K, Figure S4I-J)**. This inverse correlation was observed even on a trial-by-trial basis, as the BF-PV neural response to the reward in each trial was negatively scaled by the preceding anticipatory licking rate during the stimulus epoch ― a behavioral proxy of expectation^69,70^ **(Figure 4L, Figure S4K)**.

In summary, these results reveal that BF-PV neurons respond to behaviorally salient events such as predictive cues, rewards, punishments, and surprises, exhibiting three distinct features: (i) a selective response to behaviorally salient events, (ii) cue-related activity modulated by outcome expectation, and (iii) outcome-related activity modulated by the degree of surprise.

### A computational model of motivational salience-guided sustained attention

To interpret the observed diversity in response profiles of BF-PV neurons related to sustained attention **(Figure 2-3)** and salient events **(Figure 4)**, we employed a computational model. First, we sought to account for those three observed features of BF-PV responses to cues and outcomes described above **(Figure 4)**. A reinforcement learning (RL) framework is a natural choice to explain cue and outcome-related responses^71^. We used a temporal difference learning algorithm, TD(λ), to simulate the “Cued-Valence” and “Cued-Probability” tasks. This model uses reinforcement history to compute two key variables: the value of states, *V*, and the prediction error, i.e., the difference between expected and received value, δ, which is used to update the state values. Given that BF-PV neurons respond to cues and outcomes for both reward and punishment, that is, their signal is valence-free **(Figure 4)**, we considered only the unsigned magnitudes of these variables, |δ| and |V|. Through iterative updates across trials, the model learns the appropriate state values and develops responses to outcome-predicting cues while ignoring the task-irrelevant ones. We observed that both |δ| and |V| adjust proportionally to the reward expectations during the cue epoch, while during the outcome epoch only |δ| inversely correlates with outcome surprise (**Figure 5A-B**; top and middle rows, c.f. **Figure 4I, K**). Conversely, we observed that only |V| shows sustained elevation following the cue, aligning with the sustained activity of BF-PV neurons during the cue epoch **(Figure 4K)**. Note that this sustained cue response decayed much slower than the reward responses, indicating that it cannot be solely explained by the slow dynamics of the calcium sensor **(Figure 4K)**.

**Figure 5.**
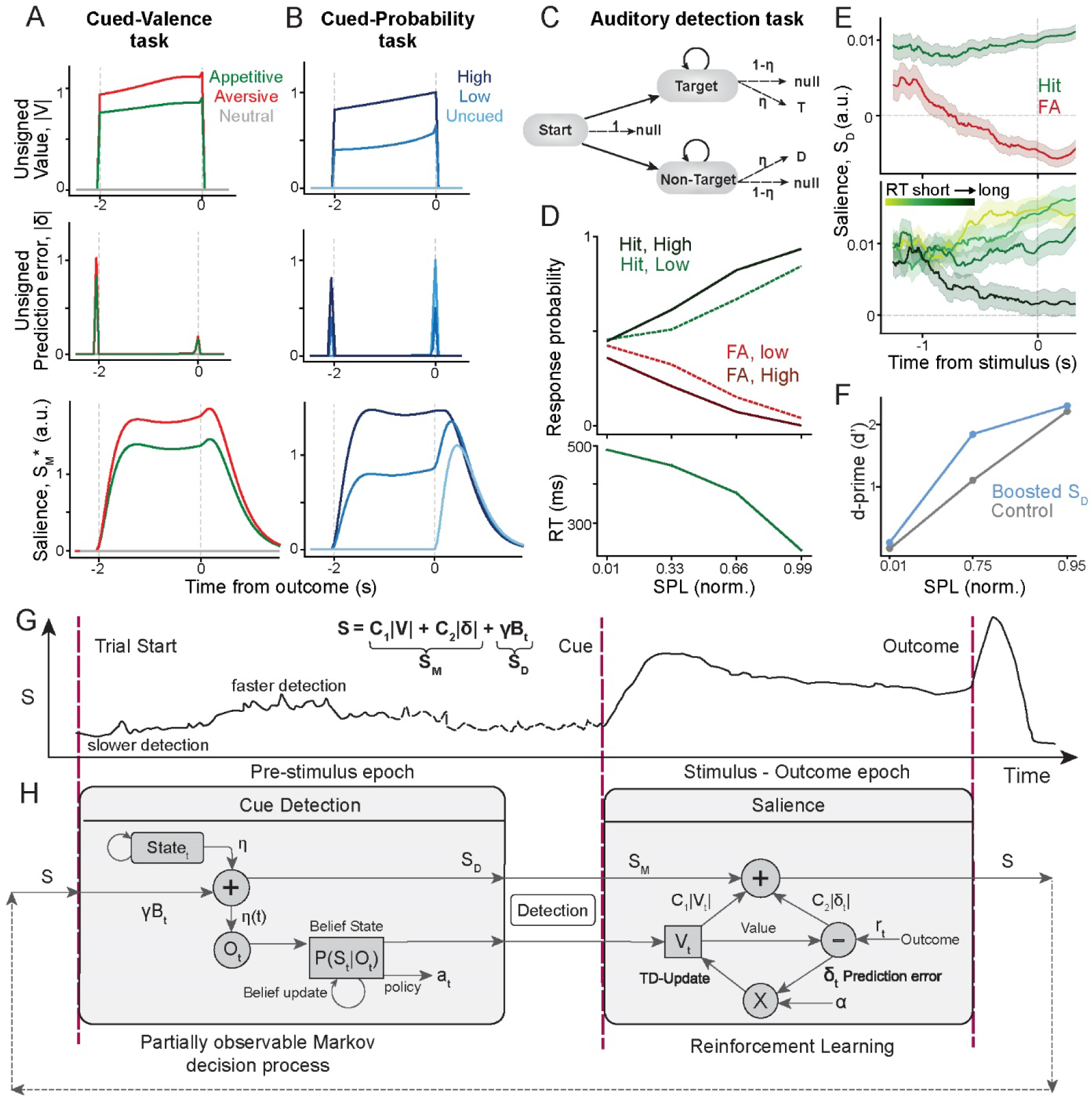
A motivational salience computation accounts for BF-PV activity. (A-B) Reinforcement learning (RL) model to simulate the Cued-Valence (A) and Cued-Probability (B) tasks. Unsigned Value, |V| (top row), and Outcome prediction error, |δ| (middle row) are depicted, aligned to the outcomes, across different trial types. The bottom row depicts the motivational salience variable, *S*_*M*_*, which qualitatively reproduces BF-PV activity during the cue-outcome epoch in Figure 4I, K. (C) Partially observed Markov decision process (POMDP) to model the auditory detection task. (D) Top, model predicted performance measured through Hit and False alarm rates, separated for trials with high (solid lines) and low (dashed lines) levels of pre-stimulus detection probabilities (median split), as a function of detection difficulty (SPL, normalized sound pressure level). Bottom, the model predicted reaction time against SPL. (E) The salience signal, *S*_*D*_ modeled as the pre-stimulus fluctuation in detection probability separated based on behavior outcomes (top) or reaction time duration (bottom). The model qualitatively reproduces BF-PV pre-stimulus activity in Figure 3A and Figure 2G, L. (F) Model predicted discriminability (d’) against detection difficulty for trials with (blue) and without (gray) boosting *S*_*D*_. (G-H) Schematic representation for the unified computational model for the salience signal, *S* representing BF-PV activity. The red, vertical dashed lines represent the onsets of salient task events: trial start, cue, and outcome. (G) A cartoon trace for a trial divided into two distinct task epochs: the foreperiod epoch on the left, and cue and outcome epochs on the right. (H) Left, in the foreperiod epoch, *S*_*D*_ represents the agent’s attentional state and determines the cue detection likelihood. The model maintains a belief state, P(s_t_|o_1_,…,o_t_) that is updated every moment based on incoming evidence. Right, in the cue and outcome epochs, *S*_*M*_ integrates responses to learn the salience of motivationally significant events – cue and outcome in a valence-free manner, while completely ignoring the task-irrelevant events.

These observations suggested that BF-PV responses could be explained by a combination of both features ― |δ| and |V| ― that are central to the concept of motivational salience. Motivational salience is the capacity of an event to drive learning and decisions in behaviors. In a reinforcement learning framework, this capacity is determined by the estimated value of state ― *V*, and the level of surprise ― δ, independent of valence (unsigned magnitude)^72–74^. Thus, we formalized “motivational salience”, *S*_*M*_, as the sum of these variables: *S*_*M*_ = *c*_1_|*V*| + *c*_2_|δ|. This model was tested by simulating the calcium transient ― as a convolution of *S*_*M*_ with calcium dynamics kernel and fitting the coefficients *c*_1_ and *c*_2_ – to match the average BF-PV responses for both the “Cued-Valence” and “Cued-Probability” tasks. The resulting adjusted salience signal, *S*_*M*_ ^∗^ demonstrates key qualitative characteristics consistent with BF-PV response tuning to cues and outcomes (**Figure 5A-B**, bottom row; cf. **Figure 4I, K**, respectively). This confirms that our model of motivational salience effectively explains the diverse BF-PV responses through a straightforward principle.

Next, we also sought to account for the observed sustained-attention-predictive activity during the pre-stimulus epoch in the auditory detection task. For this task, detecting the signal from the background noise requires a statistical inference process. Therefore, we used a partially observed Markov decision process (POMDP)^44,75^ that continually updates its belief state during the foreperiod to either “target” or “non-target” based on incoming tone stimuli **(Figure 5C)**. This model qualitatively captures changes in detection performance **(Figure 5D** top; cf. **Figure 3C)** and reaction time **(Figure 5D** bottom; cf. **Figure S2B)** as a function of stimulus strength. We sought to explain how BF-PV pre-stimulus activity predicts accuracy and reaction time in the detection task. This pre-stimulus activity is characterized by low amplitude fluctuations (**Figure 2G, 2L, 3A**), modeled as a brown noise signal, *S*_*D*_, that modulates tone detection probability. We observed that, on average, a higher value of *S*_*D*_ ― interpreted as higher attention ― leads to increased accuracy **(Figure 5D** top; cf. **Figure 3C**) and vice-versa. Additionally, when separated by trial outcomes, we observed that on average the Hit trials were preceded by a larger value of *S*_*D*_ than the False alarm outcomes (**Figure 5E** top c.f. **Figure 3A**). Moreover, a larger value of *S*_*D*_ predicts a faster reaction time in a graded fashion **(Figure 5E** bottom; cf. **Figure 2G & L)**, akin to what is expected of increased attentional modulation. We conclude that our attentional variable, *S*_*D*_, qualitatively reproduces the key characteristics displayed by BF-PV pre-stimulus activity in predicting accuracy and reaction time.

Finally, we used simulations to determine how changing *S*_*D*_ could impact detection behavior. By introducing a small, baseline increase in *S*_*D*_ in each trial, we observed that accuracy is significantly boosted for difficult-to-detect stimuli (intermediate SPL=0.75), while the effect is less obvious for easy (SPL=0.01) or hard (SPL=0.95) stimuli (**Figure 5F**). This is because at low signal-to-noise ratios (i.e. SPL=0.01) the detection of an imperceptible signal cannot be improved by enhancing the attentional state. On the other hand, in the high signal-to-noise ratio regime (i.e. SPL=0.95), the performance is not limited by the perceived stimulus strength, although it is still limited by other types of errors. Hence, the maximum behavioral effect of modulating *S*_*D*_ is expected to occur in the middle range for stimuli of moderate detectability.

In summary, our models devised two key variables, *S*_*M*_ and *S*_*D*_, that together capture the key characteristics shown by BF-PV activity across both cued-outcome and detection tasks (**Figure 5G, H)**. The key variable *S*_*M*_ represents a combination of unsigned value and outcome prediction error, which aligns with the operational definition of motivational salience. A statistical inference process that incorporates activity fluctuations as the variable *S*_*D*_ could explain detection task performance and reaction time, pointing to a computation of sustained attention.

### Elevated BF-PV activity improves sustained attention-directed behaviors

Next, we tested the causal role of BF-PV neurons in regulating behavioral performance in the detection task. Specifically, the model predicts the largest increase in accuracy at a median level of signal-to-noise ratio **(Figure 5F)**. To test this, we virally expressed a Cre-dependent channelrhodopsin (ChR2) in posterior BF-PV neurons of PV-Cre mice and implanted optic fibers above the infected areas bilaterally **(Figure 6A-B)**. Once mice learned the auditory detection task, they received bilateral laser stimulation (473nm, 0.5-2mW, 10ms, 10Hz) starting 0.5s before tone onset until outcome onset in random trials, mimicking the pre-stimulus ramped increase of BF-PV activity. We found optogenetic activation of BF-PV neurons significantly enhanced animals’ Hit rates to the “target” tone without obviously affecting False alarm rates to the “non-target” tone **(Figure 6C-D)**, thus promoting the detection discriminability **(Figure 6E).** Notably, the improvement of detection discriminability was most robust at the median level of stimulus detection difficulty **(Figure 6F)**, which matches the model prediction **(Figure 5F)**. Optogenetic activation of BF-PV neurons also decreased reaction time **(Figure 6G)**. In contrast, we observed no obvious changes in licking probability either before tone onset **(Figure 6H)**, or when animals were sated at session ends **(Figure 6I)**, suggesting no influence on general motor actions. These results demonstrate that optogenetically activating BF-PV neurons improves animals’ accuracy and response speed, replicating the effects of enhanced attentional modulation.

**Figure 6.**
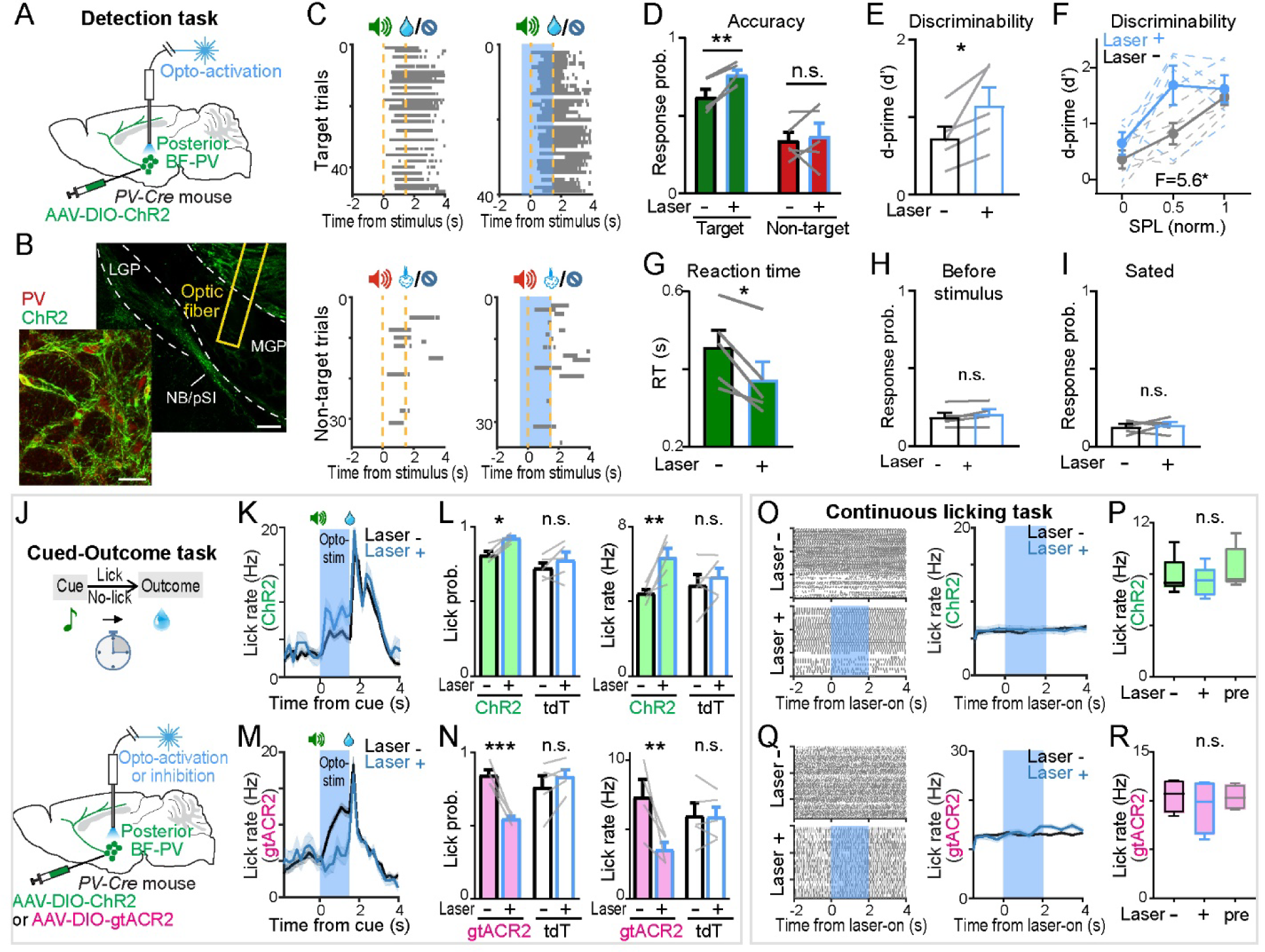
Optogenetic manipulation of BF-PV neurons regulates accuracy, reaction time, and motivated behaviors. (A-I) Optogenetic activation of BF-PV neurons in mice performing auditory detection task (n = 5 mice). (A-B) Schematic of the approach (A) and the expression of ChR2 (EYFP-tagged, green) on the membrane surface of BF-PV neurons (anti-PV staining, red, left) as well as optic fiber placement (right) in posterior BF (B). Scale bar, left, 50μm; right, 200μm. (C-D) Optogenetic activation of BF-PV neurons increases the Hit rate, but not the False alarm rate. Single session licking raster (C) and lick response probability (D) for “target” [Hit rate, p = 0.006] and “non-target” tones (False alarm rate, p = 0.70) with or without laser stimulation across animals. Blue shadowed areas in (C) indicate the window of laser stimulation starting 0.5s before stimulus onset until outcome onset (both indicated by orange lines). (E-F) Detection discriminability shown as d-prime score in trials with (Laser+, blue) or without (Laser-, black) laser stimulation [(E), p = 0.04], and separated by sound pressure level [(F), Laser+ vs Laser-, F(1, 24) = 5.6, p = 0.026; SPL, F(2, 24) = 11.6, p = 0.0003; Laser+ vs Laser-at 40db, p = 0.034]. (G-I) Optogenetic activation of BF-PV neurons decreases reaction time [(G), p = 0.02], but has no impact on lick response probability before stimulus onset [(H), p = 0.12], or after animals were sated [(I), p = 0.68]. (J-N) Optogenetic activation (K-L) or inhibition (M-N) of posterior BF-PV neurons in mice performing Pavlovian Cued-Outcome task. (J) Schematic of trial structure (top) and approach (bottom). (K, M) Average lick rate in a ChR2-(K) or gtACR2-expressing (M) mouse with or without laser stimulation. The blue area indicates the window of laser stimulation covers the cue epoch. (L) Anticipatory lick probability (left) and lick rate (right) during cue epoch are increased by laser stimulation in ChR2-expressing mice, but not in tdTomato-expressing control mice [licking probability, ChR2, p = 0.01, n = 5; tdTomato, p = 0.22, n = 5; licking rate, ChR2, p = 0.004; tdTomato, p = 0.55]. (N) Anticipatory lick probability (left) and lick rate (right) during cue epoch are decreased by laser stimulation in gtACR2-expressing mice, but not in tdTomato-expressing control mice [licking probability, gtACR2, p = 0.0005, n = 4; tdTomato, p = 0.19, n = 5; licking rate, gtACR2, p = 0.009; tdTomato, p = 0.99]. (O-R) Optogenetic activation (O-P) or inhibition (Q-R) of posterior BF-PV neurons has no impact on licking behaviors in mice doing the continuous licking task. (O, Q) Lick raster (left) and lick rate time series (right) in a ChR2-(O) or gtACR2-expressing (Q) mouse with or without laser stimulation. The blue area indicates the 2s window of laser stimulation. (P, R) Average lick rate in the 2s window without (-), with (+), or before (pre) optogenetic activation (P, [F(2, 12) = 0.7, p = 0.52, n = 5]) or optogenetic inhibition (R, [F(2, 9) = 0.3, p = 0.74, n = 4]) across animals. Data are represented as means ± SEM. *p < 0.05, **p < 0.01, ***p < 0.001, n.s., not significant; using paired t-tests for sample pairs, one-way ANOVA followed by Tukey’s multiple comparisons for groups of three or more samples, and repeated measures two-way ANOVA followed by Sidak’s multiple comparisons for groups on two independent variables. See also Figure S7.

A second model prediction is that the impacts of BF-PV activation could boost motivation specifically for cues that carry salience (e.g. task-relevant cues), as opposed to generic engagement or arousal. To test that, we virally expressed Cre-dependent light-gated neuronal activator ChR2, inhibitor gtACR2^76^, or tdTomato (as a control) in posterior BF-PV neurons bilaterally in different cohorts of PV-Cre mice, and assessed the influence of optogenetic manipulation on motivated licking behaviors toward reward-predictive cue in a Pavlovian “Cued-Outcome” task **(Figure 6J)**. Optogenetic activation of ChR2-expressing BF-PV neurons (473nm, 2-4.5mW, 10ms, 10Hz) during the cue epoch (from cue delivery until outcome onset) significantly promoted animals’ anticipatory licking probability and licking rate for the reward-predictive cue **(Figure 6K-L)**, while optogenetic inhibition of gtACR2-expressing BF-PV neurons (473nm, 0.25-0.5mW, continuous stimulation) suppressed such anticipatory licking behaviors **(Figure 6M-N)**. In contrast, the tdTomato-expressing mice were not affected by the optical stimulation **(Figure 6L, N)**. Interestingly, the optogenetic manipulation did not affect the ongoing licking behaviors in either ChR2- or gtACR2-expressing mice doing a “Continuous licking” task, where no cue was presented and every lick triggered a water delivery **(Figure 6O-R)**. These findings show that optogenetic manipulation of BF-PV activity bi-directionally regulates animals’ motivated behaviors in a task-specific manner, without influencing general motor actions.

### BF-PV neurons preferentially target cortical inhibitory neurons and produce disinhibition in the cortex

Lastly, we explored the circuit mechanism by which BF-PV neurons improve performance. A hallmark of attentional processes is to prioritize behaviorally relevant sensory information through gain modulation in the cortex^1,5,20,21^. Since the cortex receives direct projections from the BF-PV neurons as described above (**Figure 1**), we first considered which cortical cell types are the targets of BF-PV neurons. To establish this, we took a recently developed anterograde viral strategy using modified herpes simplex virus (HSV)^77^. This HSV reagent is engineered so that it can selectively replicate in Cre-positive starter neurons in the presence of Cre-dependent helper AAV, and thus transmit across a synapse into post-synaptic partner neurons in an unbiased manner. We first injected a Cre-dependent helper virus AAV-DIO-coUL6-p2a-mCherry into the BF of PV-Cre mice, then infected these neurons with the recombinant HSV-LSL-TK-GFP ΔUL6 bilaterally. As a control for potential local non-transsynaptic spread, the helper AAV was omitted in one hemisphere **(Figure 7A, Figure S5A-B)**. HSV-labeled post-synaptic neurons were found in the cortex ipsilateral to the site of helper AAV injection, with none in the contra-lateral cortex, suggesting the spread of HSV from the BF to the cortex relies on the presence of helper AAV **(pBF-PV, Figure 7B; aBF-PV, Figure S5C)**. Interestingly, the majority of the labeled post-synaptic neurons in the cortex were PV-positive inhibitory neurons (pBF-PV, 76%; aBF-PV, 63%), with a minority of CaMKII-positive principal neurons (pBF-PV, 16%; aBF-PV, 26%) **(Figure 7C-D, Figure S5D)**. The ratio of cortical PV inhibitory neurons being labeled was far above a random chance (approximately 10%^78^), representing a selectivity for targeting inhibitory neurons.

**Figure 7.**
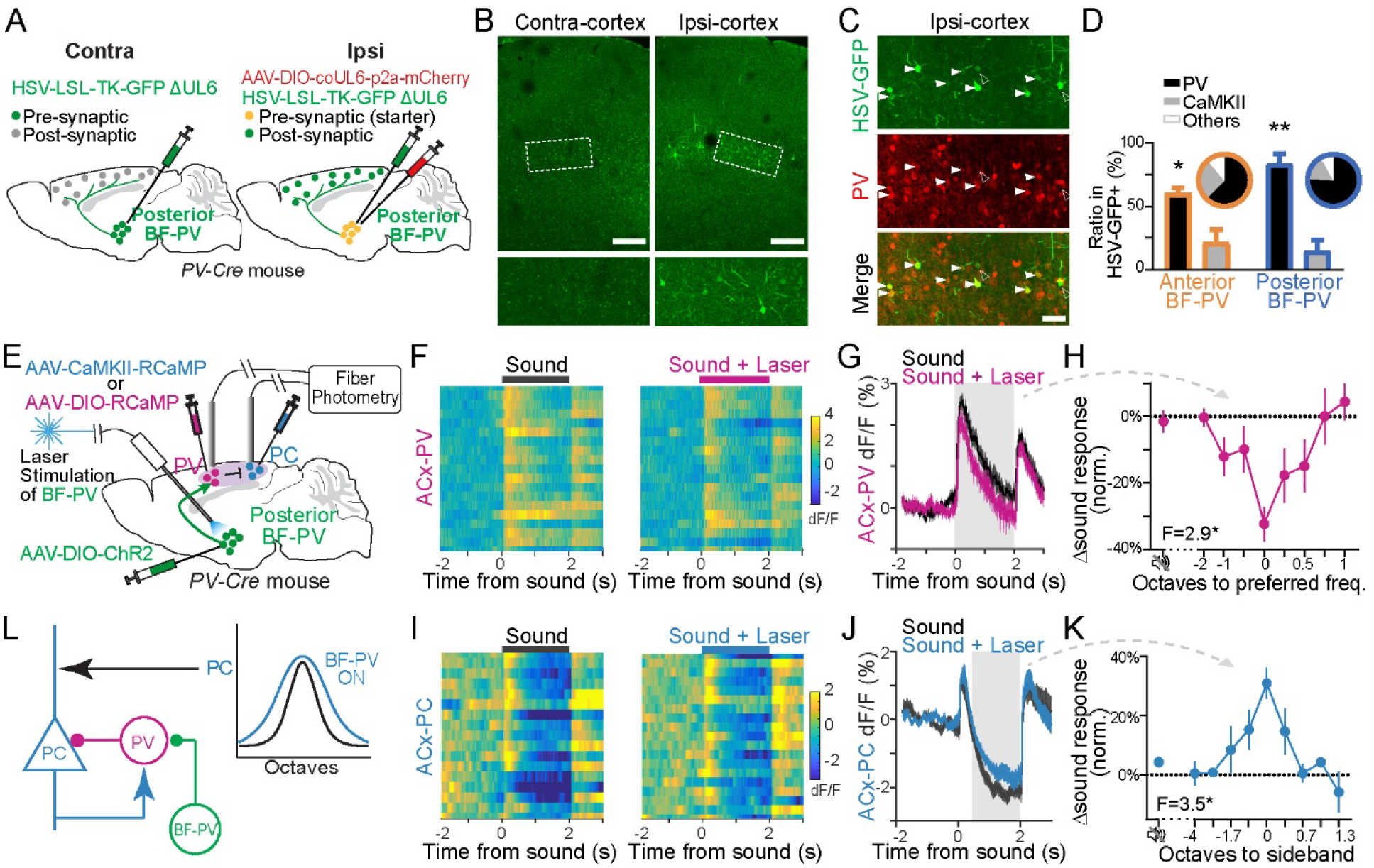
BF-PV neurons preferentially target cortical PV inhibitory neurons and generate gain modulation of auditory responses in cortex through disinhibition. (A) Schematic of anterograde trans-synaptic HSV tracing strategy to map the cortical cell types targeted by posterior BF-PV neurons. (B) HSV-GFP labeled post-synaptic neurons are found in the cortex ipsilateral to the helper AAV injection site, but not in the contralateral cortex. Bottom: enlarged images of the marked areas. Scale bar, 200μm. (C) HSV-GFP labeled post-synaptic neurons in the cortex are co-localized with PV-positive inhibitory neurons (solid arrowheads), rather than non-PV neurons (open arrowheads). Scale bar, 50μm. (D) The majority of the labeled post-synaptic neurons in the cortex are PV-positive inhibitory neurons, with a minority of CaMKII-positive principal neurons. The bar graphs compare the percentage of PV+ versus CaMKII+ cells among HSV-GFP+ cortical neurons tracing from posterior BF-PV [n = 3, p = 0.005] and anterior BF-PV [n = 3, p = 0.03]. The pie charts show the average cell-type composition of HSV-GFP+ cortical neurons tracing from posterior BF-PV [PV+, 76% (222/293); CaMKII+, 16% (25/154)] and tracing from anterior BF-PV neurons [PV+, 63% (20/32); CaMKII+, 26% (5/19)]. (E) Approach schematic for simultaneous optogenetic stimulation of posterior BF-PV neurons and fiber photometry recordings of cortical PV (F-H) and principal (I-K) neurons. (F-G) Optogenetic activation of BF-PV neurons suppresses sound responses of auditory cortex PV neurons (ACx-PV) as shown by example photometry raster (F) and average activity time series (G). The bars and the shadow areas indicate the window of sound and laser delivery. (H) The suppression effect is strongest at the preferred frequency. The suppression size at different octaves relative to the preferred frequency is measured as the difference in sound responses with versus without laser stimulation and normalized to the peak response at the preferred frequency in each animal [n = 4 mice, F(7, 22) = 2.9, p = 0.03]. (I-J) Optogenetic activation of BF-PV neurons attenuates sideband suppression (disinhibition effect) in auditory cortex principal neurons (ACx-PC) as shown by example photometry raster (I) and average activity time series (J). The sound responses are further divided into crests and troughs (shadow area). (K) The disinhibition effect is strongest at the sideband frequency with deepest troughs. The disinhibition size at different octaves relative to the sideband frequency is measured as the difference in sound responses with versus without laser stimulation and normalized to the peak response at the preferred frequency in each animal [n = 3 mice, F(8, 16) = 3.5, p = 0.02]. (L) Schematic for a hypothesized circuit mechanism of how BF-PV neurons generate gain modulation of auditory responses by inhibiting cortical PV neurons and disinhibiting cortical principal neurons. Data are represented as means ± SEM. *p < 0.05, **p < 0.01, ***p < 0.001, using unpaired two-tailed t-test for sample pairs, one-way ANOVA for groups of three or more samples. See also Figure S5, S6 & S7.

To corroborate these HSV tracing results, we used another virus AAV1, which also has the ability to travel in anterograde transsynaptic manner^79^. Although the cell types of both starter and post-synaptic cells are limited in a Cre-dependent way, it is ideal for testing connections from PV (BF) to PV (cortex) in this case. As expected, by injecting AAV1-FLEX-EGFP into the BF of PV-Cre mice, we observed post-synaptic GFP signals shown in cortical PV neurons **(Figure S5E-F)**. Together with the HSV tracing results, these findings demonstrate that BF-PV neurons preferentially target cortical inhibitory PV neurons.

Next, we tested how BF-PV activation modulates cortical activity. Because BF-PV projections are inhibitory and predominantly target cortical inhibitory interneurons, we surmised that their activation would suppress cortical inhibitory neurons and produce disinhibition. To test this, we set up simultaneous optogenetic stimulation and fiber photometry recording in awake animals. We expressed the red-shifted calcium sensor RCaMP^80^ in auditory cortex PV neurons or principal neurons to monitor their activity via fiber photometry. We also expressed ChR2^81^ in posterior BF-PV neurons for optogenetic activation **(Figure 7E, Figure S6, E)**. This allowed us to monitor the influence of BF-PV activation on cortical neural responses to sensory stimuli.

We began by establishing the baseline responses of auditory cortex PV neurons to tones of varying frequencies without optogenetic interference. The auditory cortex PV neurons were activated by both tone onset and offset, with few spontaneous activity in the quiet intervals before sound delivery **(Figure 7F**, left). Although measured at a population level, cortical PV neurons exhibited responses tuned to different tone frequencies **(Figure S6B)**, consistent with previously reported single-cell recordings^82,83^. After establishing the baseline, we investigated whether the sound responses of auditory PV neurons could be regulated by BF-PV neurons. We found optogenetic activation of BF-PV neurons during sound presentation suppressed the sound responses of cortical PV neurons when compared to the interleaved control trials **(Figure 7F-G)**. This suppressive effect was more evident for the tones producing larger sound responses (**Figure S6C**), most robust at the preferred frequency (with the largest peak of sound responses), and diminished for tones away from the preferred frequency **(Figure 7H)**. Importantly, in the absence of sound, opto-stimulation produced no discernible effects **(Figure 7H)**. These findings suggest an activity-dependent suppression of auditory PV responses by BF-PV neurons.

We then investigated whether BF-PV neurons can modulate the sound responses of auditory principal neurons. At the baseline level without optogenetic interference, cortical principal neurons also exhibited rapid phasic activation to both tone onset and offset **(Figure 7I**, left, **Figure S6D)**, and their responses were tuned to different tone frequencies **(Figure S6F)**. Notably, we observed a pronounced suppression (trough) following the peak response (crest) to tones flanking the preferred frequency **(Figure 7J, Figure S6F)**. This suppression phenomenon can be explained by the “inhibitory sideband effect” due to inputs from nearby inhibitory neurons^84–86^, thus could be potentially affected if nearby inhibitory PV neurons were suppressed by BF-PV activation. Indeed, we found optogenetic activation of BF-PV neurons during sound presentation attenuated the sideband suppression (disinhibition) effect **(Figure 7I-J).** Across animals, this disinhibitory effect was greater for tones with deeper troughs **(Figure S6G)**, and most pronounced for sidebands flanking the preferred frequency **(Figure 7K)**. Furthermore, no significant effect of opto-stimulation was observed in the absence of sound **(Figure 7K)**, suggesting that BF-PV cortical projections do not directly interrupt the activity of cortical principal cells. Instead, they seem to function as nuanced modulators of local cortical circuits, with their influence being contingent on the prevailing activity levels.

Taken together, our anatomical and physiological results show that BF-PV projections primarily target cortical inhibitory neurons to produce fast disinhibitory modulation of auditory principal neurons. This disinhibitory process regulates the contrast between the preferred frequency and sidebands, which could influence the gating of receptive field edges and reshape auditory tuning^85^. Such modulation is found to be activity-dependent, suggesting an adaptive gain modulation of sensory responses ― a putative circuit mechanism for attentional control^19,87^.

## DISCUSSION

Here we identified the circuit, behavioral, and computational functions of cortex-projecting basal forebrain PV-expressing neurons in the context of sustained attention. First, we demonstrated that BF-PV neurons send extensive and topographically organized projections to the cortex, allowing rapid and selective modulation of cortical activity **(Figure 1)**. Second, we showed that in a sustained attention task, BF-PV pre-stimulus activity (both spiking and axonal activity) tracks momentary attention fluctuations on a trial-by-trial basis (**Figure 2-3**) and their optogenetic activation improves detection performance **(Figure 6)**. Furthermore, these neurons also respond to motivationally significant events: predictive cues, rewards, and punishments, reflecting unsigned value and surprise **(Figure 4)**. This can be explained by their encoding of ‘motivational salience,’ providing a unifying account for BF-PV activation **(Figure 5)**. Motivational salience aligns with a previously proposed central computation for adaptively allocating attention based on goal relevance^66,67,88,89^. Lastly, we discovered that BF-PV neurons preferentially target cortical PV inhibitory neurons, leading to disinhibition of principal neurons and gain modulation of responses **(Figure 7)**, reminiscent of how attention modulates the gain of sensory responses^19,87^. Our findings link the behavioral and computational functions of BF-PV neurons to their underlying neural circuit mechanism, providing a framework for understanding their role in attentional control.

### BF-PV activity predicts and supports sustained attention performance

The neural activity features of BF-PV neurons display a strong correspondence with existing theories of attention. Posner and colleagues have described attentional processes as comprising three stages: alerting (sustained attention/alertness/readiness for impending stimuli), orienting (the selection towards a salient stimulus), and executive attention (the ability to flexibly regulate attention allocation according to task demands or conflicts)^2,13^. This theory reflects the multi-faceted nature of attention, and the measurement of its varied dimensions may require different behavioral tasks^3,90^. However, many previous human fMRI and EEG studies only use single task paradigm or particular task epochs, which lack a systematic understanding of the general and specific factors of attentional processes. In this study, by using a battery of behavioral tasks (e.g. sustained-attention–demanding detection task, two different cued-outcome tasks), we showed that one neural population, the cortically-projecting BF-PV neurons, displays multiple features supporting all three stages: 1) their pre-stimulus activity predicts behavioral markers of sustained attention ― reaction time and detection accuracy **(Figure 2-3)**, and drives fast and accurate performance **(Figure 6)**; 2) their neural responses are oriented toward task-relevant stimuli that predict crucial outcomes over irrelevant stimuli **(Figure 4A-F & H, Figure S4H)**; 3) their stimulus responses and outcome responses are flexibly acquired through learning **(Figure S4E)** and scaled by expectation and surprise **(Figure 4G & J-L, Figure S4G & I-K)**. These features suggest that BF-PV neurons support general, but not local, attentional processes, which are aligned with the signature of sustained attention as a foundational form of attention that could generalize to improve other attentional processes^22,91,92^. It will be an interesting open question in the future to evaluate BF-PV’s functional roles under other classic tasks that each measure a specialized dimension of attention (e.g. test selective attention in Dichotic listening task^93^).

We also considered whether BF-PV neuronal activity tracks arousal, which often accompanies and influences attentive behaviors. While a certain level of arousal is necessary for sustained attention, their relationship follows an inverted-U curve^17,18^. Our observations suggest that sustained attention exhibits momentary fluctuations^22–24,59^ **(Figure 2C)** and requires computations beyond arousal **(Figure S3)**, which is distinct from the pronounced association between arousal state and performance observed in less attention-demanding vigilance task^94,95^. The relationship between arousal and attention is mirrored in BF-PV neurons: their baseline activity correlates with and modulates arousal^29,48^, while their fast response properties under behavioral task structure reflect the computations of sustained attention **(Figure 2-4**).

### BF-PV neurons encode motivational salience to guide attention allocation

The salience of outcomes (rewards/punishments) derives from three principal factors: their physical intensity (physical salience), their novelty and surprise (surprise salience), and their positively/aversively motivating impact (motivational/emotional salience). Motivational salience is determined by the value component of outcomes and can be extended to associated predictive cues through learning. These components support each other; for example, value information needs to be updated when expected and received values don’t match (surprise/prediction error)^72–74^. We observed that the cue- and outcome-related activities of BF-PV neurons exhibited features consistent with motivational salience **(Figure 4, FigureS4)**. To operationalize these complex concepts and evaluate how motivational salience drives sustained attention, we devised a computational model **(Figure 5)**. The reinforcement learning framework applied in this model has been successfully used for modeling midbrain dopaminergic neuron responses^71^. We formalized motivational salience as a combination of unsigned value and outcome prediction error. The model-predicted activity matched well with BF-PV cue- and outcome-related responses across different tasks, indicating that the computational underpinnings necessary to explain the observed features are consistent with the operational definition of motivational salience **(Figure 5A-B)**. Moreover, the model simulated the fluctuations of this motivational salience signal during the foreperiod by adding a noise signal, *S*_*D*_, successfully predicting accuracy and reaction times in the detection task and mirroring the observed trends in BF-PV neurons’ pre-stimulus activity **(Figure 5C-E)**. Our model suggests that BF-PV neurons use the computation of motivational salience to guide their neural correlates of attention fluctuation.

The idea that motivational salience guides attention allocation has been proposed for a long time but with few systematic investigations^67,88,96,97^. Sustaining attention over extended periods demands a high level of cognitive effort, which should scale with an event’s motivational importance for efficient resource utilization^22,23^. Recent findings demonstrate that attention is also captured by surprising events^98^ and stimuli predicting rewarding or aversive outcomes (referred to as value-/threat-driven attentional capture)^66,89,99^, suggesting a common mechanism based on motivational salience rather than hedonic valence. Our detection task likely involves value-based attentional capture, as animals are required to detect “target” but not “non-target” tones. The observation that the False alarm rate was not modulated by stimulus detection difficulty **(Figure 2B)** suggests Correct reject could be influenced by factors other than attention, such as disengagement and miss, and thus were excluded from the analysis.

The temporal separation of cognitive variables in distinct task epochs in our simplified model aids interpretation **(Figure 5G-H)**. However, the interdependence between sustained attention and motivational salience^100^ should be recognized. Designing specialized task paradigms to delineate how altered levels of motivational salience guide sustained attention is an interesting future direction.

### BF-PV neurons provide dis-inhibitory control of cortex with topographic organization

We found that the BF-PV neurons, unlike PV neurons in adjacent structures like thalamic reticular nucleus^101,102^, globus pallidus^103,104^ or subthalamic nucleus^105,106^, project extensively to the cortex with unique anatomical features supporting sustained attention. These include: 1) broadcasting signals cortex-wide, 2) region-specific regulation through separate anterior^47,107–110^ and posterior BF-PV pools, and 3) modular organization within each pool. Posterior BF-PV neurons predominantly target lateral sensory cortices, such as the auditory and motor cortex **(Figure 1D-F**), and exhibit similar temporal profiles of axonal activity in these target cortices **(Figure 2-4, Figure S2 & S4)**, consistent with the organization of cortex-projecting non-cholinergic BF neurons into segregated or overlapping pools based on the inter-connectivity of their cortical targets^55^.

BF-PV neurons cross-talk with the prefrontal cortex, a well-known center for top-down attentional control^111,112^, primarily targeting prefrontal PV neurons **(Figure 7A-D, Figure S5)** implicated in attention^34^. The prefrontal cortex, in turn, innervates the basal forebrain, forming synaptic connections preferentially with BF-PV but not cholinergic neurons^113^. Indeed, they can be distinguished from frontal cortex-projecting cholinergic neurons^103^ and previously reported unidentified bursting neurons^114^ in their firing properties with long latency (median at 119 ms) and high firing rate (5-80 Hz and median at 13 Hz) **(Figure S4A-B)**.

Beyond projection organization, BF-PV’s disinhibitory modulation of cortical activity provides an anatomical substrate for gain modulation **(Figure 7E-K, Figure S6)**, a hallmark of attentional processes in the cortex^19,87^. Optogenetic activation of BF-PV neurons inhibited auditory PV neurons and disinhibited auditory principal neurons by modulating their sideband inhibition, paralleling cortical PV neurons’ ability to inhibit principal neurons’ sound responses on sideband frequencies^84–86^. This allows BF-PV neurons to reshape receptive fields and shift tuning towards behaviorally relevant signals, aligning with previous reports of basal forebrain-mediated cortical map reorganization^115,116^, and consistent with BF-PV’s function in boosting cortical responses reminiscent of increased attention^49,50,117^. We postulate that BF-PV neurons serve as an upstream area to the cortex, providing fast modulation of cortical activity and mediating gain modulation to selectively enhance attended signals, as demonstrated by improved task accuracy and faster reaction times during optogenetic activation in a sensory detection task (**Figure 6**).

### BF-PV to cortical PV disinhibitory circuit operates similarly to neuromodulatory systems

Modern views of neuromodulation emphasize the ability of subcortical circuits to modulate neural activity in distant regions, rather than focusing on transmitter identity. This modulation is usually mediated by unique transmitters not locally produced, such as acetylcholine, dopamine, and norepinephrine^52,118–120^. The BF-PV to cortical PV disinhibitory circuit shares key features with classical neuromodulatory systems, despite using GABA as a transmitter. First, BF-PV neurons are located in a deep brain region but project to distant cortical areas **(Figure 1)**. Second, BF-PV neurons exhibit homogeneous response profiles **(Figure 2F**, **Figure 4C)**, similar to the activity patterns observed within neuromodulatory populations. Third, BF-PV neurons modulate cortical activity indirectly through the disinhibition of local PV interneurons **(Figure 7)**, analogous to how classic neuromodulators influence their targets via specific receptors. This disinhibitory effect can only occur when local inhibition is already engaged, allowing for context-dependent modulation of cortical activity **(Figure 7)**. Moreover, BF-PV neurons encode cognitive variables similar to those represented by other neuromodulatory systems^121–124^ albeit reflecting distinct computations and response profiles **(Figure 4-5)**.

### Basal forebrain: division of labor across cell types

The basal forebrain contains two major ascending systems: PV+ GABAergic and cholinergic neurons. Although cholinergic neurons were traditionally thought to mediate attention, largely based on some initial observations that lesions of basal forebrain cholinergic neurons damaged animals’ performance in attention tasks^27,31,33^, and cholinergic transients were associated with cue detection^125,126^. Direct recordings of BF cholinergic neurons and cortical acetylcholine release revealed that tonic acetylcholine activities correlate with arousal states^29,38,39^ while phasic responses signal reinforcement surprise^40–43^. However, BF cholinergic neurons failed to track reaction time, an attentional performance marker^44^. Furthermore, optogenetic activation of BF cholinergic neurons increased both Hit and False alarm rates, contradicting a pro-attention effect^123^. We propose that cholinergic and PV neurons in the basal forebrain support reinforcement learning and sustained attention respectively, to guide cognitive behaviors.

Intriguingly, cholinergic and PV neurons in the basal forebrain encode similar cognitive variables central to motivational salience computation but express them differently, likely controlling distinct facets of cortical function. This resonates with the historical division of attention into “attention for learning,” essential for acquiring new information, and “attention for performance,” crucial for executing familiar tasks^127^. Cholinergic encoding of unsigned prediction errors aligns with “attention for learning” theories, controlling the learning rate for associating cues with outcomes^40,41,115,128,129^. In contrast, BF-PV encoding of sustained attention aligns with “attention for performance” theories, setting the stage for biased processing of motivationally salient cues and outcomes.

The division of labor between cholinergic and PV neurons in the basal forebrain might be due to differences in their spike timing, downstream targets, and local connectivity^42,44,46^. Cholinergic neurons exhibit fast, phasic responses to reinforcement signals **(Figure S4A-B)**, likely read out by cortical VIP neurons to produce disinhibition of principal neurons’ dendrites^62,130^. In contrast, BF-PV neurons show slow, tonic, and sustained responses **(Figure 4B-C, Figure S4A)**, primarily targeting cortical PV neurons to drive the disinhibition of principal neurons’ perisomatic domains **(Figure 6-7)**. Furthermore, the unidirectional excitatory connection from cholinergic to PV neurons^29,131^ may explain why sustained attention is unique to PV neurons, while reinforcement surprise is found in both populations.

### Implications for Cognitive Disorders

Together, these findings reveal that the basal forebrain is engaged in multiplex cognitive functions, albeit served by different neuron types. Dysfunction in the basal forebrain is associated with Alzheimer’s disease, dementia, and age-related cognitive declines^35–37^. By disentangling cell-type-specific contributions, our work may illuminate new circuit targets for therapeutic interventions and provide a fresh perspective on previous basal forebrain lesions and deep brain stimulation studies^132–135^.

Targeting the PV-mediated disinhibitory circuit, for instance, may prove beneficial in disorders characterized by attentional deficits^136–139^. Our findings highlight the importance of considering cell-type-specific functions within the basal forebrain when investigating the neural basis of cognitive disorders and developing novel therapeutic strategies.

## ACKNOWLEDGEMENTS

We are grateful to Dr. Florin Albeanu, Qiaojie Xiong, Amelia Christensen, Quentin Chevy, and Qian Li for discussions and comments on the manuscript. We thank Weixin Zhong for helping with animal surgeries and data collection. This work was supported by grants NKFIH K135561, NKFIH K147097, and NAP2022-I-1/2022 from the Hungarian Academy of Sciences to B.H.

## AUTHOR CONTRIBUTIONS

Conceptualization, S.-J.L. and A.K.; Methodology, S.-J.L., B.H. and A.K.; Investigation, S.-J.L. and B.H.; Model Methodology, U.G.; Software, J.F.S; Resources, K.B.F. and E.M.C.; Formal Analysis, S.-J.L. and B.H.; Funding Acquisition, A.K.; Writing – Original Draft, S.-J.L., U.G., and A.K.; Writing – Review & Editing, S.-J.L., U.G. and A.K with the inputs from B.H. and J.F.S..

## EXPERIMENTAL MODEL AND SUBJECT DETAILS

### Animals

Adult male and female wildtype (JAX: 000664) and transgenic mice in C57BL/6J background, including PV-Cre (JAX: 017320), ChAT-IRES-Cre (JAX: 006410), PV-Ai14 (PV-Cre crossed with Ai14 (JAX: 007914), PV-H2BGFP (PV-Cre crossed with Rosa26-stopflox-H2B-GFP^140^), were used for experiments at a minimum age of 2 months. Mice under water restriction were provided scheduled access to water, and their body weight was monitored daily to ensure it remained above 85% of their initial weight. All procedures involving animals were carried out under the protocol approved by Cold Spring Harbor Laboratory Institutional Animal Care and Use Committee in accordance with National Institutes of Health guidelines.

## METHOD DETAILS

### Tracing strategies

For anterograde tracing of BF-PV projections to the cortex, PV-Cre mice were subjected to stereotaxic injections of AAV5-CAG-FLEX-tdTomato (UNC Vector Core, AV4599D, 7.8E12 GC/ml) or AAV1-CAG-FLEX-EGFP (a gift from Hongkui Zeng, Addgene #51502-AAV1) into anterior BF or posterior BF regions (coordinates provided below). After 3-4 weeks, the mice were sacrificed, and the brains were processed by standard histological and imaging procedures. The resulting images were analyzed for slice registration and axon quantification.

For retrograde tracing of cortex-projecting BF-PV neurons, mice were subjected to stereotaxic injections of different retro-tracers in cortical regions including mPFC, RSG, M1, S1, and ACx. Three different strategies were employed: (1) wildtype mice were injected with red retro-beads (1/10 dilution, Lumafluor), and the cell identity of resulting retrogradely labeled neurons was confirmed by anti-PV and anti-ChAT (choline acetyltransferase) immunostaining; (2) PV-Cre mice were injected with Cre-dependent retrograde AAVrg-hSyn-DIO-EGFP (1.4E13 GC/ml, Addgene, #50457-AAVrg) or AAVrg-FLEX-tdTomato (1.2E13 GC/ml, Addgene, #28306-AAVrg), the cell identity of labeled neurons was constrained to PV neurons and re-confirmed by anti-PV immunostaining; (3) PV-Ai14 mice were injected with AAVrg-DIO-H2B-GFP (3.5E12 GC/ml, packaged by UNC Vector Core), the resulting retrograde signals were restricted in the nuclei of PV neurons and re-confirmed by co-localization with tdTomato signals. Mice were sacrificed 5 days after retro-bead injections or 3 weeks after AAV injections. Brain sections were processed using standard histological and imaging procedures. The resulting images were subjected to slice registration and cell counting analysis.

For anterograde tracing of the postsynaptic targets of BF-PV neurons, two viral strategies were implemented: (1) HSV tracing strategy^77^: HSV has a special capsid structure that facilitates its moving along axons in the anterograde direction and travels through synapses. We used a modified herpes simplex virus (HSV), H129 LSL-TK-GFP ΔUL6. The deletion of its UL6 gene, coupled with the Cre-dependent expression of its TK gene, renders the virus only replicate in and spread from Cre-positive starter neurons, then move across one synaptic connection and label downstream synaptic partners. For the injection procedures, we first introduced UL6 selectively in BF-PV neurons by injecting a Cre-dependent helper virus AAV8 nef-2N-DIO-coUL6-p2a-mCherry (4.22E13 GC/ml, a gift from Callaway lab) into anterior or posterior BF regions of PV-Cre mice unilaterally, then after 3 weeks, injected recombinant HSV H129 LSL-TK-GFP ΔUL6 (3.2E8 pfu/ml, gift from Callaway lab) in the same area bilaterally. The mice were sacrificed 7 days later, and the labeled neurons were identified using anti-PV, anti-CaMKII and anti-GABA immunostaining; (2) AAV1 tracing strategy^79^: PV-Cre mice were injected with AAV1-CAG-FLEX-EGFP in anterior BF or posterior BF, then sacrificed after 3 weeks, and the resulting labeled signals were restricted in PV neurons and co-localized with anti-PV immunostaining. Brain sections were processed using standard histological and imaging procedures. The resulting images were subjected to slice registration and cell counting analysis.

### Stereotaxic Injection and Implantation

Stereotaxic injections were performed using standard procedures as previously described^44,141^. Briefly, mice were anesthetized with isoflurane (0.5-1.5%), head-shaved, and positioned in a stereotaxic frame with their skull leveled in both Bregma-Lambda and medial-lateral axes to ensure precise targeting (David Kopf Instruments). Local anesthesia was achieved by administering subcutaneous injections of lidocaine (2%, 0.05-0.1mL). A small cranial window was drilled, and then a glass micropipette (10-20μm tip diameter) was inserted into the target brain region to deliver the reagent (tracer, virus, etc.). At each dorsal-ventral level, 150nL of reagent was injected at a speed of 30nL/min. Then the pipette was kept in place for 5 mins before slowly retracting from the brain. Lastly, the craniotomies were covered by low-viscosity silicone elastomer sealant (Kwik-Cast, World Precision Instruments). The wound was closed by suture and tissue adhesive (3M Vetbond). All surgeries were conducted under aseptic conditions, and the body temperature was maintained with a heating pad. After surgery, post-operative analgesia was maintained via subcutaneous injections of ketoprofen (0.5mg/kg). Unless otherwise stated, mice were housed for 3 weeks following surgery to allow for recovery and viral expression before histological procedures, behavioral training, or recordings.

For the tracing experiments, the following coordinates were used: anterior BF (AP: +1.3 mm, ML: 0.8 mm, DV: 5.0mm, and AP: +0.8mm, ML: 0.8mm, DV: 5.2 mm from brain surface), posterior BF (AP: -0.4mm, ML: 1.8mm, DV: 4.5mm, and AP: -0.8mm, ML: 2.1mm, DV: 4.5 and 4.0mm), mPFC (AP: +1.9mm, ML: 0.5mm, DV: 2.2 and 1.7mm), RSG (AP: -1.5mm, ML: 0.3mm, DV: 0.5mm and AP: -1.0mm, ML: 0.3mm, DV: 0.5mm), M1 (AP: +2.4mm, ML: 1.8mm, DV: 1.0mm and AP: +3.0mm, ML: 1.5mm, DV: 1.0mm), S1(AP: +1.9mm, ML: 3.0mm, DV: 0.7mm and AP: -1.5mm, ML: 3.0mm, DV: 0.7mm) and ACx (AP: -2.0mm, ML: 4.0-4.5mm, DV: 0.5mm and AP: -2.5mm, ML: 4.0-4.5mm, DV: 0.8mm).

The implantation of microdrives was conducted as previously described^44^. First, PV-Cre mice were injected with Cre-dependent AAV expressing a special version of channelrhodopsin with fast kinetics (AAV5-Ef1a-DIO-ChETA-EYFP^81^, a gift from Karl Deisseroth, UNC Vector Core, 3.9E12 GC/ml, 300nL) in the posterior BF region (AP: -0.8mm, ML: 2.0mm, DV: 4.5 and 4.0mm, 150nL at each depth), then their skull surface were coated with adhesive cement (C&V Metabond, Parkell). The tetrode bundle of microdrive was pre-coated with DiI (Life Technologies), then lowered vertically into the brain to a depth of 2-2.6mm. The microdrive was secured to the skull using UV adhesive (EM ESPE) and dental acrylic (Lang Dental). Ground and reference electrodes were implanted in the parietal cortex bilaterally, followed by the placement of a titanium head bar for head fixation. The positions of tetrode leads and recorded neurons in every animal were recovered by histology and tetrode track reconstruction after the last recording session **(Figure S7D)**.

For optic fiber implantations in the cortex, PV-Cre mice were first injected with Cre-dependent AAV expressing an axon-enriched calcium indicator GCaMP6 (AAV5-hSynapsin1-FLEx-axon-GCaMP6s, a gift from Lin Tian, Addgene, # 112010-AAV5, 1.35E13 GC/ml) in the posterior BF (AP: -0.8mm, ML: 2.0mm, DV: 4.5 and 4.0mm, 150nL at each depth), as well as AAV1-CAG-tdTomato (Addgene, #105554-AAV1, 2E11 GC/ml) in the ACx (AP: -2.0mm, ML: 4.0 or 4.5mm, DV: 0.5-0.8mm, 75nL) and M1 (AP: +2.4mm, ML: 1.8mm, DV: 1.0mm, 75nL). Then optic fibers (400um core, 0.48NA, 2.5mm zirconia ferrule, 5mm or 8mm fiber length, Doric Lenses) were inserted into the brain about 0.2mm above the target sites, and secured to the skull using UV adhesive and dental acrylic. The positions of fibers in every animal were confirmed within aim regions by histology after the last recording session **(Figure S7A-B)**.

For *in vivo* optogenetic-photometry experiment, PV-Cre mice were injected with AAV5-Ef1a-DIO-ChETA-EYFP (3.9E12 GC/ml, UNC Vector Core) in the posterior BF (AP: -0.8mm, ML: 2.0mm, DV: 4.5 and 4.0mm, 150nL at each depth), as well as AAV1-Syn-FLEX-NES-jRCaMP1a (a gift from Douglas Kim & GENIE Project, Addgene, #100848-AAV1, 4.6E12 GC/ml) or AAV9-CaMKII-RCaMP2 (1.5E12 or 5E12 GC/ml) in the ipsilateral ACx (AP: -2.0mm, ML: 4.0 or 4.5mm, DV: 0.7 and 0.5mm, 75nL at each depth). Then the fiber for optogenetic activation (200μm core, 0.39 NA) was implanted into the posterior BF and another fiber for fiber photometry (400μm core, 0.48NA) was implanted into the ACx about 0.2mm above the target site. The positions of fibers in every animal were confirmed within aim regions by histology after the last recording session **(Figure S7B-C)**.

For the optogenetic manipulation of behaviors, PV-Cre mice were injected with AAV5-Ef1a-DIO-ChR2(H134R)-EYFP (a gift from Karl Deisseroth, UNC Vector Core, 3.2E12 GC/ml), AAV1-hSyn1-SIO-stGtACR2-FusionRed (a gift from Ofer Yizhar, Addgene, # 105677-AAV1, 6E12 GC/ml), or AAV1-FLEX-tdTomato (7E12 GC/ml, Addgene, # 28306-AAV1) in the posterior BF bilaterally (AP: -0.8mm, ML: 2.0mm, DV: 4.5 and 4.0mm, 150nL at each depth), then optic fibers (200μm core, 0.39 NA) were implanted bilaterally into the brain about 0.2mm above the target sites. The positions of fibers in every animal were confirmed within aim regions by histology after the last behavior session **(Figure S7C)**.

### Histology

Histology experiments were conducted following standard procedures as previously described^141^. Briefly, brains were cut into 60μm coronal sections by a vibratome (Leica). Alternatively, brains were precipitated in a PBS solution containing 30% sucrose for 36-48 h at 4℃, then cut into 30μm coronal sections using a Cryostat (Leica). The brain sections were then incubated in a blocking solution (PBS containing 5% bovine serum albumin and 0.3% Triton X-100) for 1hr at room temperature, followed by incubation overnight at 4℃ in the primary antibody solution (blocking solution plus diluted primary antibodies). The following primary antibodies were used: Rabbit anti-PV (Thermo Fisher Scientific, Cat# PA1-933, RRID:AB_2173898, 1:500); Mouse anti-PV (Swant, Cat# 235, RRID: AB_10000343, 1:500); Goat anti-PV (Swant, Cat# PVG-213, RRID: AB_2650496, 1:500); Goat anti-ChAT (Millipore, Cat# AB144, RRID: AB_90650, 1:500); Rabbit anti-CaMKII (Abcam, Cat# 52476, RRID: AB_868641, 1:500); Mouse anti-CaMKII (Cell Signaling Technology, Cat# 50049, RRID: AB_2721906, 1:250); Rabbit anti-GABA (Sigma, Cat# A2052, RRID: AB_477652, 1:500); Rabbit anti-RFP (Rockland, Cat# 600-401-379, RRID: AB_2209751, 1:2000); Chicken anti-RFP (Rockland, Cat# 600-901-379, RRID: AB_10704808, 1:500); Rabbit anti-GFP Alexa Fluor 488-conjugated (Molecular Probes, Cat# A-21311, RRID: AB_221477, 1:500); Mouse anti-GFP (Santa Cruz Biotechnology, Cat# sc-9996, RRID: AB_627695, 1:500); Rabbit anti-Cre (Millipore, Cat# 69050-3, RRID: AB_10806983, 1:500). The next day after washing in PBS, brain sections were incubated in the secondary antibody solution (blocking solution plus diluted Alexa Fluor conjugated secondary antibodies, Invitrogen/Molecular Probes, 1:500 or Jackson ImmunoReseach Labs, 1:250) at room temperature for 2 hrs. After washing, brain sections were mounted in a glycerol mounting medium with or without DAPI as needed (Electron Microscopy Sciences, Cat# 17989-61, # 17989-51).

### Slice Imaging, Registration, and Quantification

To study the distribution of BF-PV’s cortical projections in an unbiased stereological way, one out of every six coronal sections (60μm) was selected for immunostaining and counting. For each brain, approximately 15 sections were selected to cover the majority of cortical regions, spanning about 5.4 mm in the anterior-posterior axis. To map the locations of cortex-projecting PV neurons throughout the BF, one out of every three coronal sections (60μm) was selected for immunostaining and counting, thus for each brain approximately 9 sections were selected to cover the entire BF regions, spanning from about 1.8 mm anterior to Bregma until 1.6 mm posterior to Bregma. Images were acquired using an LSM710 or LSM780 confocal laser scanning microscope (Carl Zeiss Microscopy). Slices from the same brain were always processed together with the same settings to ensure consistency in morphological analyses.

To read out the precise positions of neurons, axons, and fiber tracks in the brain, a MATLAB-based software SHARP-track was employed^53^. This involved navigating each histology image to a matching reference slice from the atlas (Allen Institute Common Coordinate Framework, CCFv3), which could be freely rotated in anterior-posterior, medial-lateral, and dorsal-ventral axes. The histological images were then geometrically transformed and projected onto the reference atlas. By registering relevant slices in this manner, all the marked neurons, axons and fiber tracks on histological images could be mapped into the atlas and plotted on a 3D brain model. This method allows for more precise alignment of brain sections that were sliced at imperfect angles, and provides automated coordinate extraction and sub-region segmentation that are particularly useful for brain regions with ambiguous borders like the BF and cortex.

Axon quantification and cell counting were conducted using Image-Pro Plus software (Media Cybernetics, RRID: SCR_007369). For each image, the region of interest was outlined from the DAPI channel, and signals within this region were threshold-processed to remove noise pixels. Clusters of continuous pixels above the threshold were identified as axons or cells, and the integrated axon signals (total areas × signal intensity) or the total cell numbers were automatically counted by the software. To validate the automated counting, manual counting was performed for some samples blinded to the experimental conditions, and the results from automated and manual counting were similar. Lastly, the measured numbers from each sub-region were normalized to the summed number in each animal.

### Behavior Setup

The animals were tested in sound-isolated and ventilated chambers that were specifically designed for the behavioral experiments. The chambers were equipped with waterspouts to deliver water rewards, an infrared lickometer (Island Motion Co.) to detect licks, LEDs for light signals, and a separate tubing system for air-puff delivery. The timing and volume of water and air flow were precisely controlled and calibrated by solenoid valves. Auditory stimuli were presented using two calibrated speakers to ensure standardized sound pressure levels. In addition to the licking detection, pupil diameter was recorded using separate video cameras (Point Grey) under the control of the Bonsai framework (Lopes et al., 2014). Stimulus delivery, behavioral monitoring, optical stimulation, photometry, and spike recording data acquisition were all controlled using a Windows computer and the open-source Bpod/PulsePal behavioral control system (Sanworks LLC, US) with custom software written in MATLAB^142^.

### Auditory Detection Task

The auditory detection task that requires high sustained attention was set up as previously described^44^. Briefly, water-restricted mice were trained in habituation sessions first, during which head-fixed mice learned to lick the water spout to get an un-cued water reward (5μL) either under operant conditioning (get water if pokes, 10-20 trials) or under direct water delivery at randomized intervals (exponential, mean = 3s, cap at 9s). Next, mice proceeded to the direct delivery phase, where they received water if they licked within 90s after a “target” tone (4 or 20 kHz, 50- or 60-dB sound pressure level, 0.5s). Typically, after one or two sessions, mice developed tone-reward association and anticipatory licking in response to tone delivery. After that, mice were introduced to the full task in which they had to lick after the “target” tone (4 or 20 kHz, 10-50dB, 0.5s) to obtain a water reward (5 or 8μL) and withhold licking after the “non-target” tone (8 or 10 kHz, 10-50dB, 0.5s) to avoid punishment (a mild facial air-puff for 200ms). Each trial was concluded with one of the four following possible outcomes: two types of correct trials, Hit (lick response to “target” tone) or Correct reject (no-lick response to “non-target” tone), and two types of incorrect trials, Miss (no-lick response to “target” tone) or False alarm (lick response to “non-target” tone). Unless otherwise stated, each trial consisted of a pre-start period (with LED-on for at least 1.5s) followed by a start signal (signaled by LED-off), a foreperiod (see below), a stimulus and response window (starts with a 0.5s tone stimulus, 0.6s or 1s in total), a delay period (see below), a short outcome delivery period (depending on the duration of water or air-puff delivery) and a post-outcome period (4s) **(Figure 4A)**. A no-lick policy was forced before the tone onset, which means the trial was restarted if mice licked during the pre-start period or foreperiod. All types of trials were randomly interleaved. Recording data was included in the analysis only after stable psychometric performance was reached. Each animal contributed at least three sessions.

To manipulate temporal attention, the foreperiod was randomly chosen either from an exponential (mean = 2.4s, cutoffs at 0.1s and 4s; or mean = 1.4s, cutoffs at 0.1s and 5s) or a bimodal distribution (mixture of two Gaussians and a uniform distribution with mixing probabilities 0.35, 0.35 and 0.3; means, 0.3 and 2 s; s.d., 0.15 s; cutoffs at 0.1 and 3 s). Exponential foreperiod distribution results in a constant hazard rate, with an equal probability of stimulus occurrence throughout the foreperiod on any given trial. Bimodal foreperiod distribution results in temporal expectation with the stimulus more likely to be delivered at two time points according to the modes, determined by the subjective hazard model^44,64^. To separate the lick response from the actual reward response, a delay period (Gaussian distribution, mean = 0.5s, cutoffs at 0.2s and 1s, applied in all animals except for 3 mice in spike recordings) was applied in between the first lick response and outcome onset.

To increase attention demands, several strategies were applied: (1) the random duration of the foreperiod introduced temporal uncertainty about the timing of stimulus onset; (2) the short response window (0.6s in spike recording or 1s in photometry recording) mandated quick detection; (3) the graded detection difficulty by introducing fainter tones (10, 20, 30, 40, 50 dB) embedded in background white noise (10-60dB, adjusted based on previous performance to keep relatively constant False alarm rates).

To analyze animals’ behavioral performance, the psychometric curve was measured as *P_Hit_* (ratio of Hits / “target” trials) and *P_FA_* (ratio of False alarms / “non-target” trials) at different sound detection difficulties. The discrimination curve was measured as d-prime (d’) score: *d’ = norminv(P_Hit_) – norminv(P_FA_),* where *norminv* is the inverse of the cumulative normal function. Values of *P_Hit_* and *P_FA_* were truncated between 0.01 and 0.99, setting the maximum d’ to 4.65.

Since reaction time (RT) varies across different sessions and animals, and rarely conforms to a normal Gaussian distribution, here RTs were normalized as reaction time percentile rank (RT*) to quantify how fast a given RT relative to other RTs in the same session. RT* was calculated as the percentage of RTs shorter than the given RT, and was computed as *RT*(t) = (L + 0.5 * E) / N * 100*, where *L* is the number of RTs in the same session that is less than *t*, *E* is the number of RTs equal to *t*, and *N* is the total number of RTs in the same session.

### Cued-Valence Task and Cued-Probability Task

The Cued-Valence task and Cued-Probability task are head-fixed auditory Pavlovian conditioning tasks, similar to those previously described^40^. Briefly, water-restricted mice underwent habituation training, followed by a simplified auditory Pavlovian conditioning training stage where a tone was paired with water reward (8μL). After developing tone-reward association and anticipatory licking in response to tone delivery, mice were introduced to either the Cued-Valence task or the Cued-Probability task. In the Cued-Valence task, three distinct tones (randomly chosen from 4, 8, 10, 15, 20kHz, 50dB) were associated with water reward (8μL), air puff (200ms) or nothing, respectively. In the Cued-Probability task, two different tones (randomly chosen from 4, 8, 10, 15, 20kHz, 50dB) were paired with water reward with a probability of 100% or 50%, respectively. Additionally, in 10% of trials, the water reward was delivered with no proceeding tone. Unless otherwise stated, each trial consisted of a 0.5s tone stimulus followed by a 1.5s delay, a short outcome delivery period, and a post-outcome period (4s). All types of trials were randomly interleaved. The inter-trial interval was randomly chosen from an exponential distribution (mean = 2s, cap at 6s). Each animal contributed at least three sessions.

### Optogenetic Experiments

#### Auditory detection task

The water-restricted mice were trained similarly as in the detection task described above, except that the interval between tone onset and outcome onset was fixed for optical stimulation. In addition, to avoid confusing the animals, the trial-start LED signal was omitted, and the patch cords and optic ferrules were covered to prevent the animals from seeing the flashing laser light. Once the mice demonstrated stable psychometric performance, they were tested under optogenetic manipulation. In 30-40% of random trials, a burst of laser pulses (λ = 473 nm, 0.5-2mW power, 10ms pulse width at 10Hz frequency) was delivered for optogenetic activation starting 0.5s before tone onset and lasting until outcome onset. As for the optogenetic inhibition during the foreperiod in a detection task, however, we found it very challenging technically. Because during the quiescent foreperiod before cue onset, the increased cortical PV activity under opto-inhibition of BF-PV produces rebound activity (not shown) in cortical pyramidal neurons^143^.

#### Cued-Outcome task

The water-restricted mice were trained in a simplified auditory Pavlovian conditioning task similar to those described above. Each trial consisted of a 0.5s tone stimulus followed by a 1s delay, and then the water reward (8μL) was delivered. Once the mice developed stable anticipatory licking in response to tone delivery, they were tested under optogenetic manipulation. In 30-40% of trials, a burst of laser pulses (λ = 473 nm) was delivered during both the tone and delay period (1.5s in total). Optogenetic activation was performed with laser pulses of 2-4.5mW power, 10ms pulse width at 10Hz frequency, while optogenetic inhibition was performed with a single continuous pulse of 0.25-0.5mW.

#### Continuous licking task

Water-restricted mice were trained in habituation sessions first, then proceeded to the continuous licking task. Each lick to the water spout triggered a delivery of 0.75μL water. Usually, it took mice one or two sessions to achieve stable continuous licking. The mice were subsequently tested under optogenetic manipulation. Each trial in the continuous licking task was further divided into a 2s pre-laser period (baseline), a 2s laser period, and a 2s post-laser period. In 20% of trials, a burst of laser pulses (λ = 473 nm) was delivered during the laser period. Optogenetic activation was performed with laser pulses of 2-4.5mW power, 10ms pulse width at 10Hz frequency, while optogenetic inhibition was performed with a single continuous pulse of 0.25-0.5mW.

#### Simultaneous opto-activation and photometry

in this experiment, the activity of cortical neurons was recorded by fiber photometry while optogenetically activating BF-PV neurons. Each trial consisted of a 2s baseline period, a 2s tone stimulus period (60dB tones with varying frequencies), and a 2s post-tone period followed by an inter-trial-interval randomly ranging from 0 to 6s (exponential distribution with mean at 2s). In 50% of trials, the tone delivery was accompanied by a burst of laser pulses (λ = 473nm, 2mW power, 1ms pulse width, 10Hz; 2s burst length). The laser-plus-sound trials and sound-only trials were alternately presented for a block of 20 trials (10 trials for each type) at a fixed sound frequency, then switched to another block with a different sound frequency (sound was randomly selected from the following frequencies: 0, 1, 2.5, 5, 10, 15, 20, 25, 30, 35, 40 kHz).

### *In Vivo* Extracellular Electrophysiological Recording

#### Extracellular tetrode recording

we employed the miniaturized microdrive that housed a bundle of 8 tetrodes and an optic fiber (50μm core) as previously described^44^, which allows for optogenetic-tagging and independent day-to-day positioning of tetrodes for long-term recordings of single neurons in the brain. Briefly, we used a DigitalLynx data acquisition system (Neuralynx) and Cheetah data acquisition software (Neuralynx) for spike data collection. A pre-amplifier (Neuralynx Headstage HS-36) and a fine wire tether (TETH-HS-36-FWT, Neuralynx) were used to link the microdrive to DigitalLynx. Continuous broadband (0.1-9000 Hz) data was collected from each tetrode referenced to an electrode placed in the parietal cortex or a silent tetrode channel. Unit signals were filtered with a bandwidth of 600-6000 Hz and digitized at 32.552 kHz. Digital input was recorded from the behavior control system at the time of stimulus presentations to synchronize neural and behavioral data.

#### Optogenetic tagging

For optogenetic identification of BF-PV neurons, bursts of brief laser pulses (λ = 473 nm, 1 ms pulse width, 5-40 Hz; 2s burst length; 3s inter-burst-interval) were delivered in the brain both before and after each behavioral session. Laser power was in the range of 0.16-16mW and adjusted as necessary to avoid light-induced photoelectric artifacts or population spikes that might mask individual action potentials. The trigger TTL pulses were also recorded by the data acquisition system for synchronization. The tetrodes and the optic fiber were advanced 20-100μm after each data acquisition session based on single unit activity and the presence or absence of light-evoked potentials.

#### Spike sorting

data analysis was carried out in MClust 3.5 software (A.D. Redish). Cluster quality was measured using isolation distance and L-ratio calculated based on two features, the full spike amplitude and the first principal component of the waveform. Putative single neurons with isolation distance > 20 and L-ratio < 0.15 were included. Autocorrelations were inspected for refractory period violations and putative units with insufficient refractory period were excluded. Spike-shape correlations between spontaneous and light-induced spikes were calculated for all BF-PV neurons. The significance of photoactivation was assessed during offline analyses by the SALT test-based spike latency distributions after light pulses, compared to a surrogate distribution using Jensen-Shannon divergence (information radius). Neurons with p < 0.01 were considered light-activated.

#### Data analysis

we calculated event-aligned peri-event time histograms (PETHs) and corresponding raster plots for individual neurons. Individual PETHs were Z-scored by the mean and standard deviation of the baseline window. The baseline window was defined as a 1.2s-long window (from -1.2s to 0s) before sound stimulus onset. Reward response latency was determined by the peaks in the PETHs aligned to the reward. To examine whether pre-stimulus firing rates of some BF-PV neurons are predictive of animals’ reaction time (RT). RTs in Hit trials were regressed against the firing rates during the 0.25s window before or after sound stimulus onset on a trial-by-trial basis (trials with RT>0.1s were excluded to avoid accidental/impulsive responses). In addition, trials were grouped as quartiles based on RT length (Quartile 1: 0-25 percentile; Quartile 2: 25-50 percentile; Quartile 3: 50-75 percentile; Quartile 4: 75-100 percentile), then PETHs and pre-stimulus firing rates (mean value in 0.25s window before stimulus onset) were calculated for different these groups and Z-scored for individual animals.

#### Tetrode track reconstruction

To trace the locations of recorded neurons, we performed electrolytic lesions, slice registration, and track reconstruction. Briefly, after the last recording session, mice were anesthetized and electrolytic lesions were made through individual tetrode leads (5-30μA for 5s; stimulus isolator, World Precision Instruments or A-M Systems). Next, mice were sacrificed and histology processes were carried out as mentioned above. The images of coronal brain slices were geometrically transformed and projected onto the reference brain atlas so that the DiI traces of the electrode tracks and the lesion site (final protruding length of tetrodes) could be mapped into the atlas. Finally, the coordinates of detected neurons can be recovered by the depth estimates based on electrode descent distance in each recording session. Since we mainly focused on the basal forebrain sub-regions enriched in auditory cortex-projecting PV neurons based on our retrograde tracing experiments, only neurons located in the following areas were included (a total of 154 well-isolated units, including 18 optogenetic-identified PV neurons): nucleus basalis, posterior substantia innominata, border area of globus pallidus and internal capsule.

### Fiber Photometry

#### Fiber photometry recordings

Fiber photometry recordings were set up as previously described^40,141^. Briefly, the excitation beam for the green channel was produced by a 490nm LED light source (M470F3 Thorlabs), then collimated via an aspheric condenser (ACL25416U Thorlabs), passed through an excitation filter (ET470/24M Chroma), bounced off a dichroic mirror (T495LPXR Chroma). The excitation beam for the red channel was produced by a 565nm LED light source (M656F3 Thorlabs), collimated via an aspheric lens (F240FC-A Thorlabs), passed through an excitation filter ET569/25X and passed through the same dichroic mirror. The excitation beams were then launched into a 400um core, 0.48NA fiber patch cable using an aspheric objective lens (A240TM-A Thorlabs). Fluorescence excitation and detection were both accomplished through one multimode optical fiber that was connected to the animal. Fluorescence emitted was passed through an emission filter (ET525/50M and ET630/75M Chroma for the green and red channel, respectively), focused with a plano-convex lens (f = 30 LA1805-A Thorlabs), and collected using an amplified photodiode (IM 2151 New Focus). To ensure the proper separation of the two channels, the fluorescence signal was amplitude-modulated by sinusoidally varying the command voltage of the LED driver (LEDD1B Thorlabs) with two different frequencies (531 Hz and 211 Hz) and demodulated before data processing^144,145^. Data were acquired using a data acquisition card (PCIe-6321 National Instruments) and synchronized with the behavioral task using Bpod and custom software written in MATLAB.

#### Fluorescence signal processing

Fluorescence signals were expressed as the relative fluorescence *Z score = (dF(t)-F_mean_) / F_SD_*, where *dF(t)* is the *(F-F_0_)* value at time *t*, and *F_0_* is the mean fluorescence value for a given trial across the baseline window, *F_mean_* and *F_SD_* are the mean and standard deviation of the *dF* values over the baseline window averaged across all trials within a given session. The baseline window was defined as a 4s-long window (from -4s to 0s) before stimulus onset by default. For those experiments to evaluate the sustained differences preceding stimulus onset, the baseline window was brought further ahead starting from -4s before stimulus onset.

To carefully control for the possibility that the photometry signal might show neural activity-independent fluctuations, e.g. due to movement or auto-fluorescence, we simultaneously recorded tdTomato fluorescence in the red channel. The fluorescence of tdTomato did not reveal tonic and phasic modulations as shown by GCaMP6s signals, suggesting that the GCaMP6s signals of BF-PV axons that we had observed could not be explained by neural activity-independent fluctuations **(Figure S2F)**.

#### Photometry data analysis

Due to the prolonged decay of fluorescence signal using GCamP6s, the neural response to a given event was calculated trial-by-trial as the relative increase of the peak fluorescence signal (value at 90 percentile during event epoch) compared to the pre-event signal (in a 250ms window before event onset). The stimulus/CS epoch was defined as the window between tone onset to outcome onset. The outcome/US epoch was defined as a 1s window after outcome onset **(Figure 6D)**.

To examine whether BF-PV axonal activity recorded by fiber photometry is predictive of RT, we first calculated the cross-correlation between RTs in Hit trials and photometry fluorescence signals in each 0.1s bin during a 2s window (20 bins) before stimulus onset. The most robust negative correlation was found in time bins located from -0.5s to 0s before stimulus onset (**Figure 2L & Figure S2I**). Thus we selected this 0.5s pre-stimulus window to evaluate how well BF-PV activity can predict upcoming RT across all animals. Trials were grouped as quartiles based on pre-stimulus activity (Quartile 1: 0-25 percentile; Quartile 2: 25-50 percentile; Quartile 3: 50-75 percentile; Quartile 4: 75-100 percentile), then in each group the average RTs were calculated.

We also tested whether BF-PV axonal activity before stimulus onset predicts animals’ performance. Trials were grouped as low- and high-activity trials (median split) or as quartiles based on pre-stimulus activity level, then in each group the psychometric curve and discrimination curve were calculated.

#### Pupil Monitoring

The monitoring and analysis of pupil diameter fluctuations in behaving animals were conducted as previously described^40^. Briefly, a separate video camera (Point Grey) under the control of the Bonsai framework was used to record the pupil dilation of one eye. An infrared light was used to provide oblique illumination to the eye. In the meantime, a blue LED with adjusted brightness was placed close to the opposite eye to induce moderate pupillary contraction. Pupil diameter was calculated using a custom MATLAB program. After manually choosing regions of interest (ROI) and selecting a threshold to isolate the eye area and pupil area, a morphological dilation/erosion algorithm was used to automatically consolidate the pupil region. A linear regression was used to calculate the diameter of the best-fit circle to the perimeter of the pupil region. Total pixel intensity within the eye ROI was extracted for analysis of blink response. The value of pupil diameter during blink response was calculated as NaN and excluded from quantification. For the normalization of eye area or pupil diameter, the traces were normalized as *dP/P = (P(t) – P_0_)/P_0_*, where *P(t)* is the value at time *t, P_0_*is the mean value for a given trial across the baseline window (from -4s to 0s before stimulus onset). To test whether pupil diameter is predictive of behavioral performance or BF-PV activity, trials were grouped as small- and large-pupil trials (median split) or as quartiles based on the pupil diameter during the 0.5s pre-stimulus window, then in each group, the average RT percentile rank, d-prime, and BF-PV activity level were calculated.

### Model

### Partially observable Markov decision process for the auditory detection task

We simulated the auditory detection tasks using a partially observable Markov decision process^44,75^ to simulate how the accumulated likelihood of detecting the tone changes based on stimulus detection difficulty. This task model consisted of three states: ‘start’, ‘target’, and ‘non-target’. Each trial starts in the ‘start’ state and leaving the ‘start’ state was governed by the conditional probability of a stimulus to occur in the present time bin provided no stimulus had been presented before. In other words, the transition probability from ‘start’ to other states, denoted by q(t), was determined by the hazard function of stimulus onset, resulting in an exponential foreperiod distribution (with the same parameters as for the mice). The ‘target’ and ‘non-target’ states were assigned equal probabilities, mimicking the detection task learned by the mice. The ‘start’ state generated no output observations. The ‘target’ state generated a T observation with η probability and no observation with 1-η probability, reflecting the stochastic detection of the tone stimuli. Symmetrically, the ‘non-target’ state generated a D output with η probability and no output with 1-η probability. The detection probability, η, depended on the stimulus difficulty – defined as the cumulative probability of detecting at least one observation during the tone period – set to 0.01, 0.33, 0.66, and 0.99 for four levels of difficulty. The time step was set to 0.01 s. During each trial, mice only had access to the cumulative outputs without direct access to the hidden states. Therefore, they had to make decisions about the stimuli based on the ambiguous observations by performing Bayesian inference:

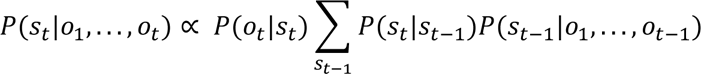

Where *s*_*t*_ ∈ {′*start*′, ′*target*′, ′*non* − *target*′} is the hidden state at time-step t, *o*_*t*_ ∈ {*null, T, D*} is the observation made by the agent at t, *P*(*s*_*t*_|*s*_*t*−1_) refers to the transition probability between states (as determined by the task) from time t-1 to t, *P*(*o*_*t*_ |*s*_*t*_) is the probability of making an observation *o*_*t*_ in the hidden state *s*_*t*_ and finally, *P*(*s*_*t*_ |*o*_1_, …, *o*_*t*_) is the inferred posterior probability after making t observations. At each time step, the agent updates the posterior using a hidden Markov model (HMM). Since trials of varying difficulty were randomly interleaved in the task without providing mice with explicit information about difficulty, an average level of detection probability was used for the inference. Response (lick) probability at each time step was defined by a combination of the calculated posterior probabilities as,

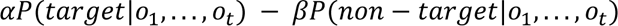

where α and β were constant parameters set to 0.05 and 0.03, respectively. If a lick occurred within 1s of stimulus onset (tone duration in the detection task), a response was assigned to the trial. Responses in the ‘target’ state resulted in Hits and responses in the ‘non-target’ state were counted as False alarms. The lack of responses in these states produced Misses and Correct rejects, respectively. Reaction time was determined by the time from tone onset to the animal’s response. Psychometric functions and reaction times produced by the HMM closely reproduced those from the auditory detection task.

**Table 1:** State transition matrix.

|  | *start* | *target* | *non-target* |
| --- | --- | --- | --- |
| *start* | $1 - q(t)$ | $q(t)/2$ | $q(t)/2$ |
| *target* | 0 | 1 | 0 |
| *non-target* | 0 | 0 | 1 |

Further, we modeled the pre-stimulus activity, with fluctuations driven by an auto-correlated noise signal. This noise influences the animal’s immediate stimulus detection probability,

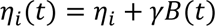

where η_*i*_ ∈ {0.01,0.33, 0.66,0.99} is the baseline, cumulative detection probability for stimulus in trial type *i*, B(t) is Brownian noise scaled by γ(= 0.1). In this formulation, the first term, η_*i*_ represents the task difficulty (set externally through the trial type), whereas the second term represents the impact of fluctuating attention on the animal’s cue detection, and consequently on the task outcome. Due to the introduced time pressure of the detection task, the animal, at various time points, could falsely perceive a tone. This is especially likely at low stimulus strengths. To model this, we also introduced a constant, false-positive rate, *f* in the detection likelihood. The cumulative false-positive rate was set to about 0.4. Note, that conversion from a cumulative (at least detected once in the tone period) to a point detection probability (at each time step) requires a geometric transform. To perform the causal manipulation (**Figure 4F**), we introduced a baseline increase in the detection likelihood (Δη∼0.15, cumulative) for each simulated stimulus strength.

**Table 2:** Observation matrix for each state.

|  | *null* | *T* | *D* |
| --- | --- | --- | --- |
| *start* | $1 - 2f$ | $f$ | $f$ |
| *target* | $1 - \eta - 2f$ | $\eta + f$ | $f$ |
| *non-target* | $1 - \eta - 2f$ | $f$ | $\eta + f$ |

Enhanced pre-stimulus activity is associated with increased accuracy, consistent with observed trends in BF-PV neurons’ pre-stimulus activity.

### Reinforcement learning for the Cued-Outcome tasks

We simulated the Cued-Outcome tasks using a TD(λ) reinforcement learning algorithm and captured how the value, *V(t)*, and outcome prediction error (OPE), δ(*t*) vary during a trial. Across all trial types, each trial is simulated for 3*s*, with the cue at time 0.5*s* and reward delivery at time 2.5*s*. The state space was modeled using a complete serial compound (CSC), where the trial is tiled with 25*ms* of serial different states. All weights were initialized with 0. In each trial, eligibility traces for all the states were initialized at 0. At each time step, V is given by

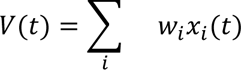

where w represents the learned weights for each state. In the CSC representation, the cue at time t, x(t) is a vector of signals {*x*_1_(*t*), …, *x*_2_(*t*)} that represents the cue for variable lengths of time into the future, such that, *x*_*i*_ (*t*) is 1 exactly *i* time-steps after the presentation of the cue in the trial and 0 otherwise^146,147^. Similarly, OPE is calculated by

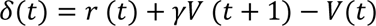

where r is the trial outcome as an arbitrary scalar (+1 for a water reward and -1.25 for the air-puff punishment). Also, the eligibility traces were updated as

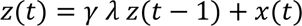

where *z*(*t*) is eligibility traces for the state localized at time t; γ is a discount factor that measures the importance of future rewards; λ is the decay parameter for the eligibility traces. Ultimately, the learned weights are updated using the following rule

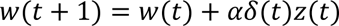

where α is the learning rate.

Once the variables, V and OPE, are simulated, we linearly combine them into the operationalized motivational salience variable as *cc*_1_|*V*(*t*)| + *c*_2_|δ(*t*)|, where the |·| is the modulus operator, and the parameters were fitted to approximately, *c*_1_ = 0.08; *c*_2_ = 0.92. Finally, to mimic the dynamics of the GCaMP signals recorded by photometry, the motivational salience variable was convolved with a representative г;-distribution trace (α = 3; β = 0.225). We used the following parameters across the Cued-Outcome tasks: discount factor, γ = 0.995; λ = 0.75; learning rate, α = 0.1, time step, Δ*t* = 0.025*t*. For the “Cued-Valence” tasks, we also introduced a very small number of trials (∼2.5%), for each trial type, where no outcome was delivered (reward or punishment).

### Quantification and statistical analysis

Statistical analyses were performed using MATLAB or commercial software (GraphPad Prism; GraphPad Software, CA). Statistical significance was tested using paired t-tests for sample pairs, one-way ANOVA for groups of three or more samples followed by Tukey’s multiple comparisons, and repeated measures two-way ANOVA for groups on two independent variables followed by Sidak’s multiple comparisons unless otherwise specified. Data are presented as means ± SEM. *p < 0.05, **p < 0.01, ***p < 0.001, n.s., not significant.

## SUPPLEMENTARY FIGURES AND LEGENDS

**Figure S1.**
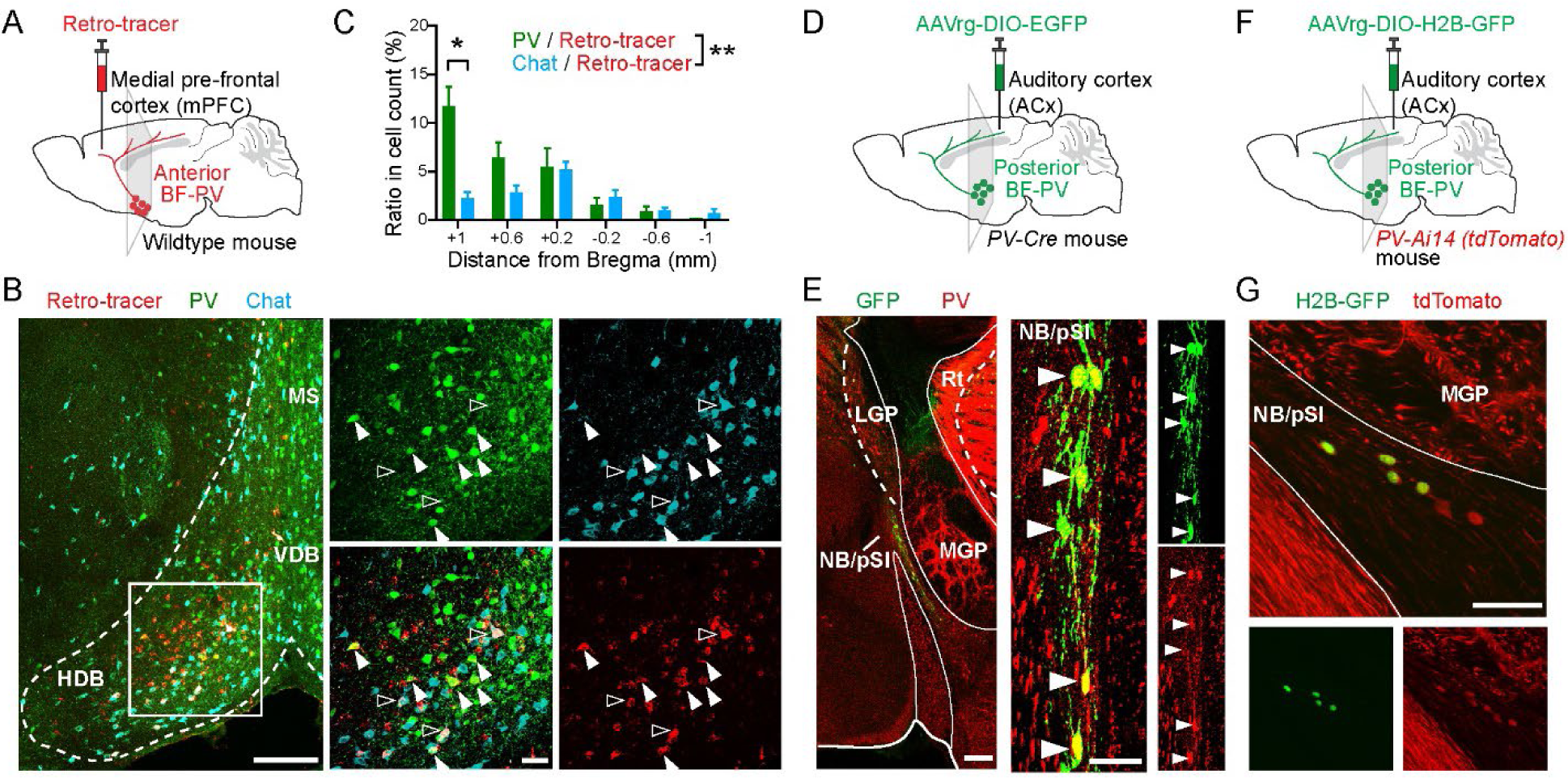
BF-PV neurons show topographic projections to the cortex, related to Figure 1. (A-C) Mapping the locations of PV and cholinergic neurons that target medial cortical regions. (A) Schematic of the retrograde tracing approach. (B) Retro-tracer injection in the medial prefrontal cortex (mPFC) labels PV and cholinergic neurons in the anterior basal forebrain. Enlarged images of the boxed area from the left are shown on the right. Solid arrowheads, mPFC-projecting BF-PV neurons labeled by co-localization of retrograde signal (red) and anti-PV immunostaining signal (green). Open arrowheads, mPFC-projecting cholinergic neurons labeled by co-localization of retrograde signal (red) and anti-Chat immunostaining signal (cyan). MS, medial septum nucleus; VDB, vertical diagonal band; HDB, horizontal diagonal band. Scale bar, left: 200μm, right: 50μm. (C) Cortex-projecting BF-PV and cholinergic neurons display different distribution patterns. The distribution of labeled mPFC/RSC-projecting PV+ and Chat+ cells is shown in each anterior-posterior (AP) coronal plane in individual mice and averaged across animals [Interaction between cell-types and AP-position, F(5, 40) = 4.9, P = 0.001; PV vs Chat at AP +1mm, p = 0.016. n = 8 mice for PV cell count, n = 3 (out of those 8 mice) for Chat cell count, using mixed-model two-way ANOVA followed by Sidak’s multiple comparison tests]. (D-E) Injection of Cre-dependent retrograde AAV expressing GFP into the auditory cortex in PV-Cre mouse (D) labels PV neurons in the posterior basal forebrain. Arrowheads, ACx-projecting BF-PV neurons labeled by co-localization of retrograde GFP (green) and anti-PV immunostaining signal (red) (E). NB, nucleus basalis; pSI, posterior substantia innominata; LGP, lateral globus pallidus; MGP, medial globus pallidus; Rt, reticular thalamic nucleus. Scale bar, left: 200μm, right: 50μm. (F-G) Injection of Cre-dependent retrograde AAV expressing nuclear-localized GFP (H2B-GFP) into the auditory cortex in PV-Ai14 mouse (PV-Cre crossed with Ai14 reporter line result in tdTomato expression in all the PV neurons) (F) labels PV neurons in the posterior basal forebrain. ACx-projecting BF-PV neurons are labeled by co-localization of retrograde GFP (green) and tdTomato (red) (G). Scale bar, 200μm.

**Figure S2.**
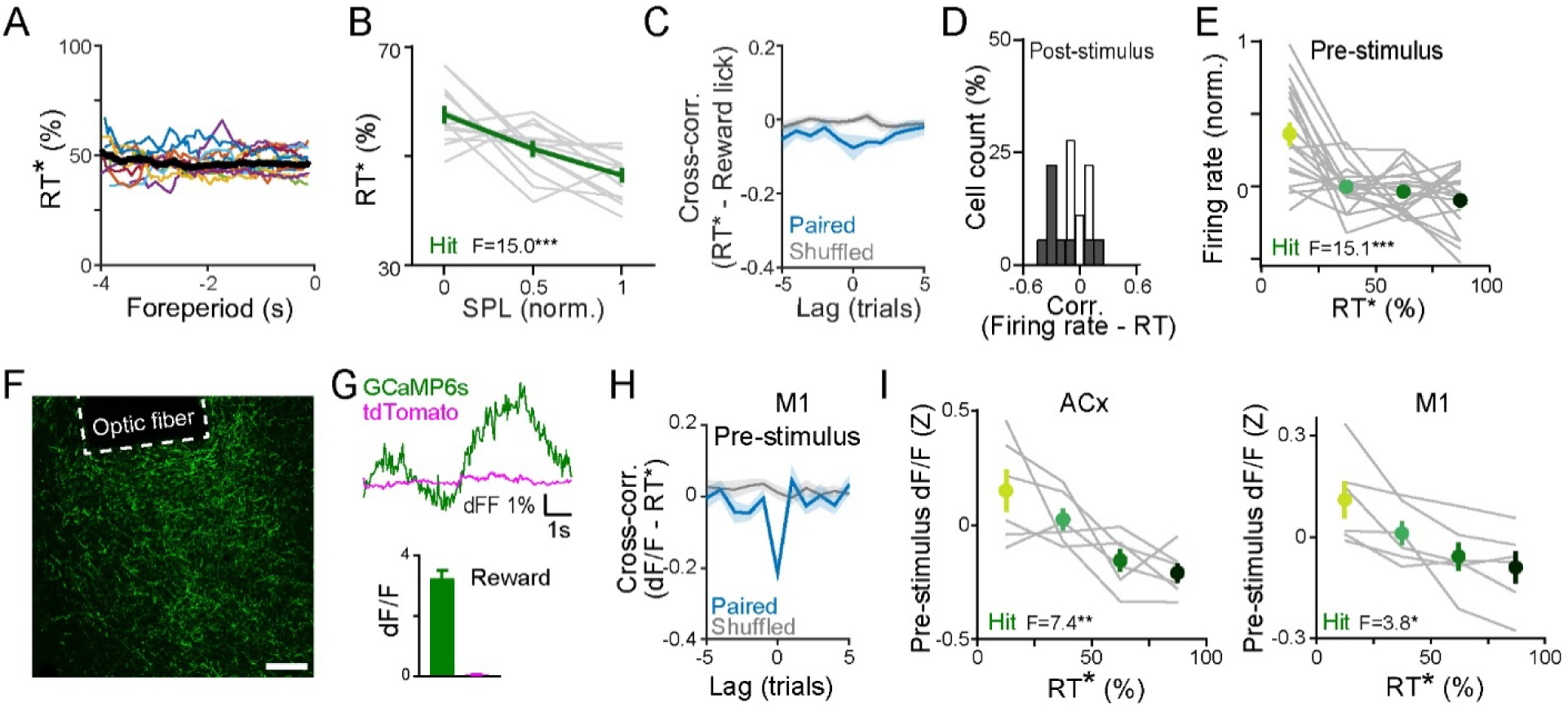
BF-PV’s axonal activity in the auditory and motor cortex predicts reaction time and accuracy, related to Figure 2. (A-C) Behavioral analysis for mice in the sustained attention task (n = 12 mice). (A) The foreperiod duration had no obvious impact on mice’s reaction times. Traces of reaction time (rank, sliding window, 5 trials) were plotted as a function of foreperiod duration for individual mice (thin curves) and pooled across animals (thick black curve). (B) Average reaction time (rank) as a function of stimulus detection difficulty for individual mice (gray curve) and grand average (green curve) [F(2, 33) = 15.0, P < 0.0001; SPL0.5, P = 0.01, SPL1, P < 0.0001, compared to SPL0]. (C) Cross-correlation between reaction time and reward licking rate shows a weak correlation. (D) Distribution of the correlation coefficient between post-stimulus firing rate and reaction time for individual neurons [skewness = -0.12]. The shaded represent neurons with significant correlation [p < 0.05, Pearson]. (E) Population spiking activities correlate with reaction time. Average pre-stimulus firing rates in Hit trials grouped by reaction time duration in individual neurons (gray) and grand average (green) [F(3, 68) = 15.1, P < 0.0001]. (F) Representative image showing the placement of optic fiber above axon-GCaMP6s-positive BF-PV fibers (green) in cortex. Scale bar, 100μm. (G) Reference tdTomato channel shows no correlation with axon-GCaMP6s signals or specific task events. Top, example axon-GCaMP6s trace (green) along with a simultaneously recorded reference tdTomato trace (magenta) acquired during behavior. Scale bars, 1% of dF/F fluorescence change, and 1s of time. Bottom, comparison of the response magnitude in reward epoch between axon-GCaMP6s versus tdTomato channel. (H) Same as in Figure 2K, except that data is from the motor cortex (n = 6). (I) Pre-stimulus axonal activities correlate with reaction time. Average pre-stimulus activity of BF-PV’s axons in the auditory cortex (G) [F(3, 20) = 7.4, p = 0.002]) and in the motor cortex (K) [F(3, 20) = 3.8, p = 0.03]) grouped by reaction time duration in individual mice (gray) and grand average (green). Data are represented as means ± SEM. *p < 0.05, **p < 0.01, ***p < 0.001, n.s., not significant; using repeated measures one-way ANOVA followed by Tukey’s multiple comparisons for groups of three or more samples, unless otherwise specified.

**Figure S3.**
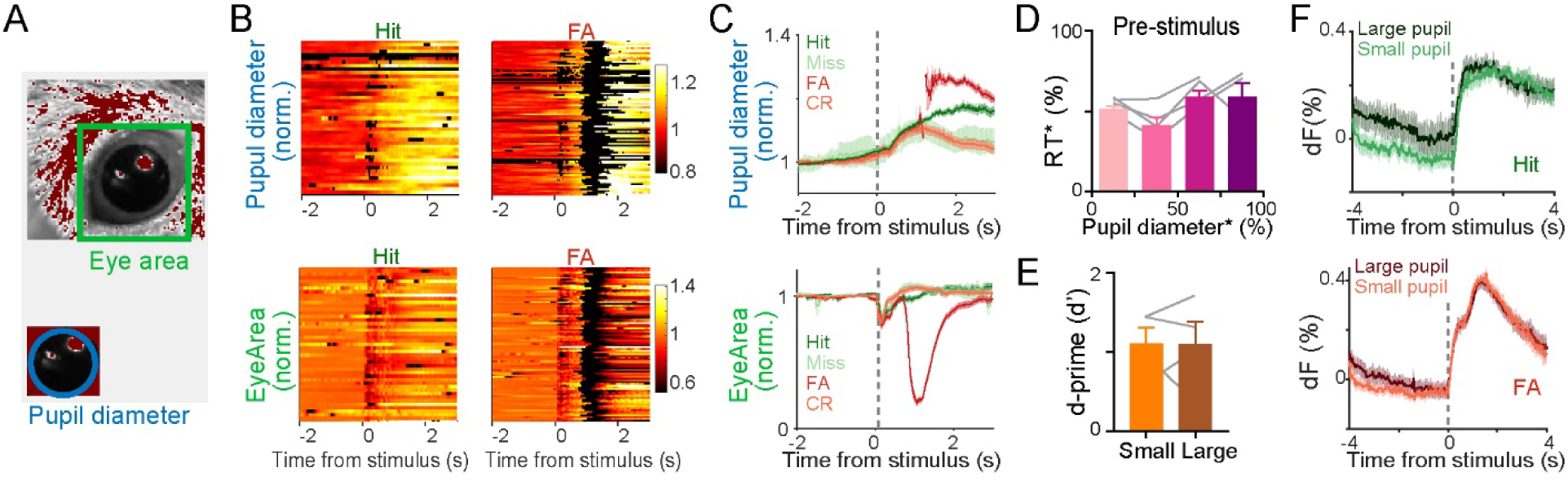
Arousal level does not predict reaction time, accuracy, or BF-PV activity, related to Figure 3. (A) Schematic of eye area and pupil diameter measurement. (B-C) Representative raster (B) and average plots (C) of pupil diameter (top) and eye area (bottom) aligned to stimulus onset in trials with different outcomes. Note that some trials display severe disruption of pupil diameter detection (black) caused by factors like eye blinking, and thus were excluded from average plots. (D-E) Average reaction time (rank) in Hit trials (D) and detection discriminability (E, d-prime score) are measured as a function of pre-stimulus pupil diameter (in a 500ms window preceding stimulus onset, quartile split or medium split) for individual mice (gray curves) and grand average (colored). Neither reaction time [F(3, 11) = 3.1, p = 0.07, n = 4] or detection discriminability [p = 0.98, n = 4] is systematically associated with pre-stimulus pupil diameter. (F) Average BF-PV axonal activity in the auditory cortex aligned to stimulus onset in Hit (top) and False alarm (bottom) trials grouped by large or small (medium split) pupil diameter during the baseline period (4s-long window before stimulus onset). Data are represented as means ± SEM. *p < 0.05, **p < 0.01, ***p < 0.001, n.s., not significant; using paired t-tests for sample pairs, one-way ANOVA followed by Tukey’s multiple comparisons for groups of three or more samples.

**Figure S4.**
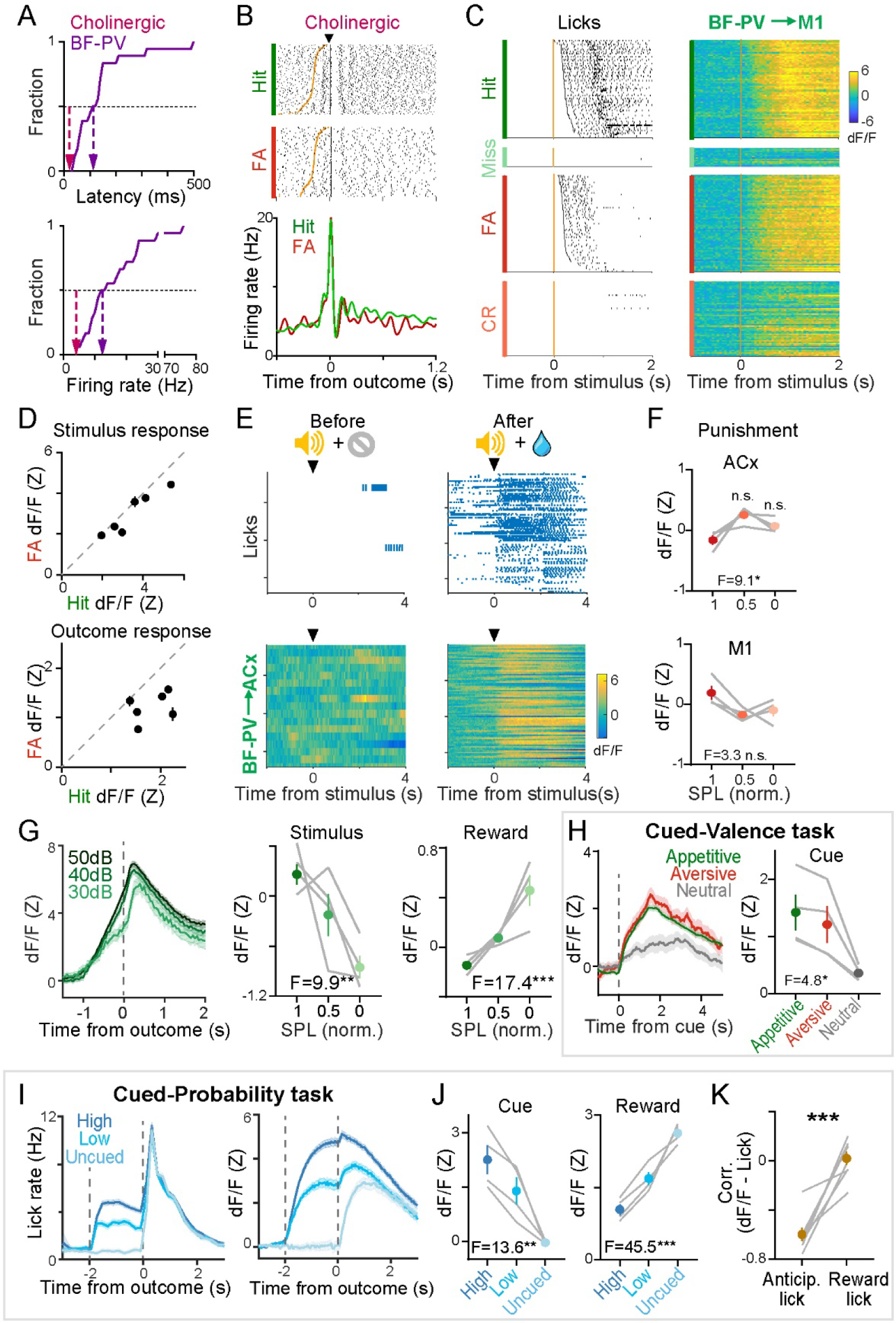
BF-PV’s axonal activity in the motor cortex also grades with unsigned value and outcome surprise, related to Figure 4. (A) Cumulative histogram of the reward response latency (top) and baseline firing rate (bottom) of all identified BF-PV neurons. The median values are indicated by purple arrows, and compared with that of cholinergic neurons (pink arrows, adapted from Hangya et al., 2015). (B) Same as Figure 4B, except that data are from an identified cholinergic neuron in the posterior basal forebrain showing fast and phasic response to reward and punishment. (C) Raster plots of licks and BF-PV axonal activity in the motor cortex (M1) for different trial types from an example animal performing the auditory detection task. Trials are aligned to stimulus onset and sorted by reaction time (orange lines: stimulus onset). (D) Same as Figure 4F, except that data are from M1 [stimulus response, p = 0.06; outcome response, p = 0.01; n = 6]. (E) Raster plots of licks (top) and BF-PV axonal activity in the auditory cortex (ACx, bottom) before and after a sound stimulus was paired with upcoming water reward. (F) Same as Figure 4G, except that data are the responses to punishment in False alarm trials recorded from BF-PV’s axons in ACx and M1. Response magnitude is normalized to mean punishment responses within subject and shown in individual mice (gray curves) and grand average (red) [ACx, F(1.2, 3.6) = 9.1, p = 0.042; SPL 50dB vs 40dB, p = 0.10; SPL 50dB vs 30dB, p = 0.09, n = 4. M1, F(1.0, 3.1) = 3.3, p = 0.16, n = 4. Using repeated measures one-way ANOVA followed by Tukey’s multiple-comparison tests]. (G) Same as Figure 4G, except that data are from M1 [Stimulus response, F(2, 9) = 9.9, p = 0.005; Reward response, F(2, 9) = 17.4, p = 0.0008; n = 4]. (H) Same as Figure 4I, except that data are from M1 [F(2, 9) = 4.8, p = 0.04]. (I-K) Same as Figure 4K-L, except that data are from lick rate and M1 [Cue, F(2, 9) = 13.6, p = 0.002; Reward, F(2, 9) = 45.5, p < 0.0001 in (J); p = 0.0002 in (K)]. Data are represented as means ± SEM. *p < 0.05, **p < 0.01, ***p < 0.001, n.s., not significant; using paired t-tests for sample pairs, one-way ANOVA followed by Tukey’s multiple comparisons for groups of three or more samples, unless otherwise specified.

**Figure S5.**
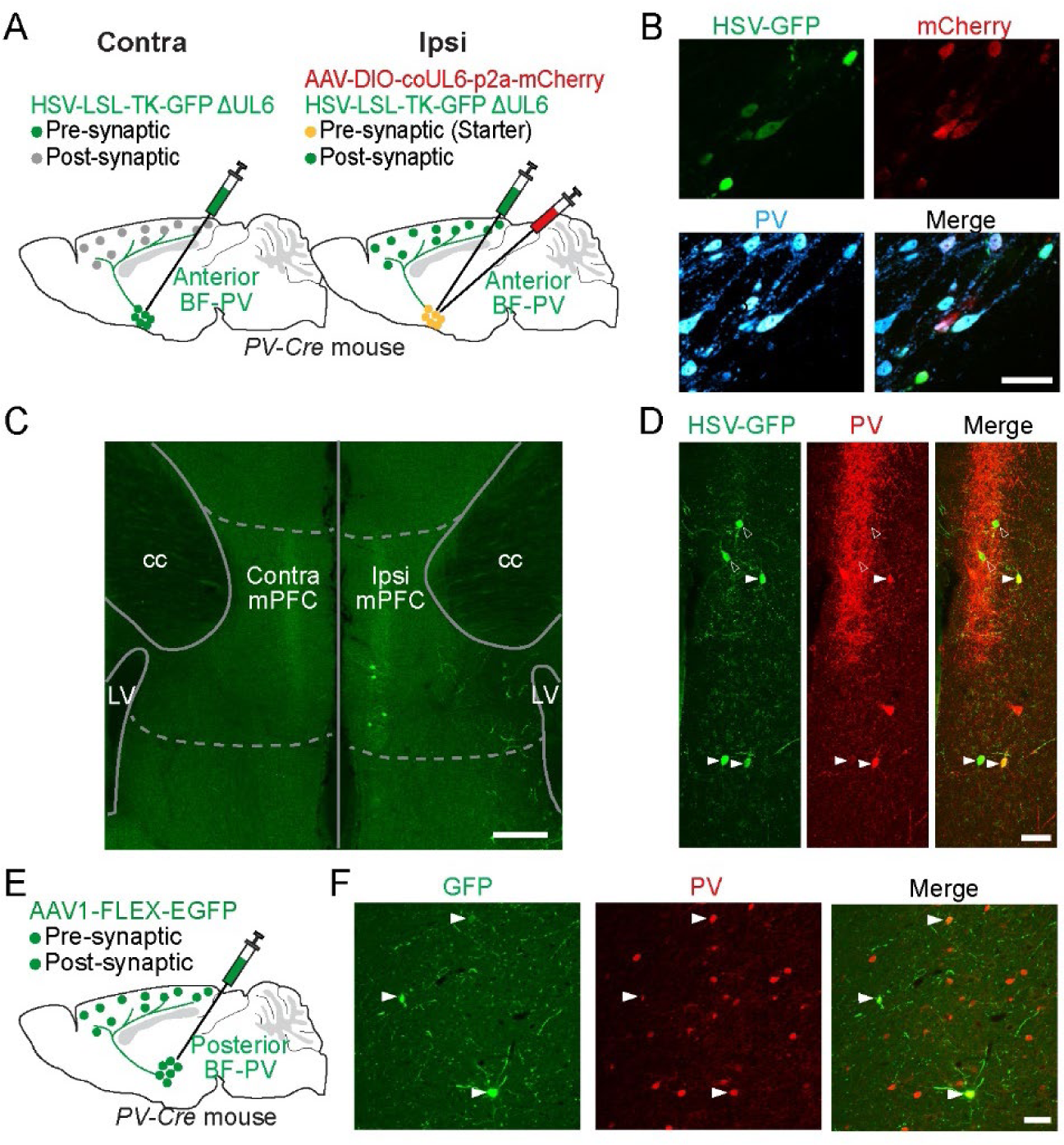
Anterior BF-PV neurons also primarily target cortical PV neurons, related to Figure 7. (A) Schematic of HSV tracing strategy to map the cortical cell types targeted by anterior BF-PV neurons. (B) Helper AAV (mCherry+, red) and HSV (GFP+, green) specifically infect PV neurons (PV+, cyan) at the injection site. Scale bar, 50μm. (C-D) Same as Figure 7B-C, except that the starter cells are anterior BF-PV neurons, and the labeled post-synaptic neurons are shown in the medial prefrontal cortex. (E-F) Anterograde trans-synaptic AAV1 tracing approach (E) labels cortical PV neurons (F, solid arrowheads) as the post-synaptic partner of BF-PV neurons. Scale bar, 50μm.

**Figure S6.**
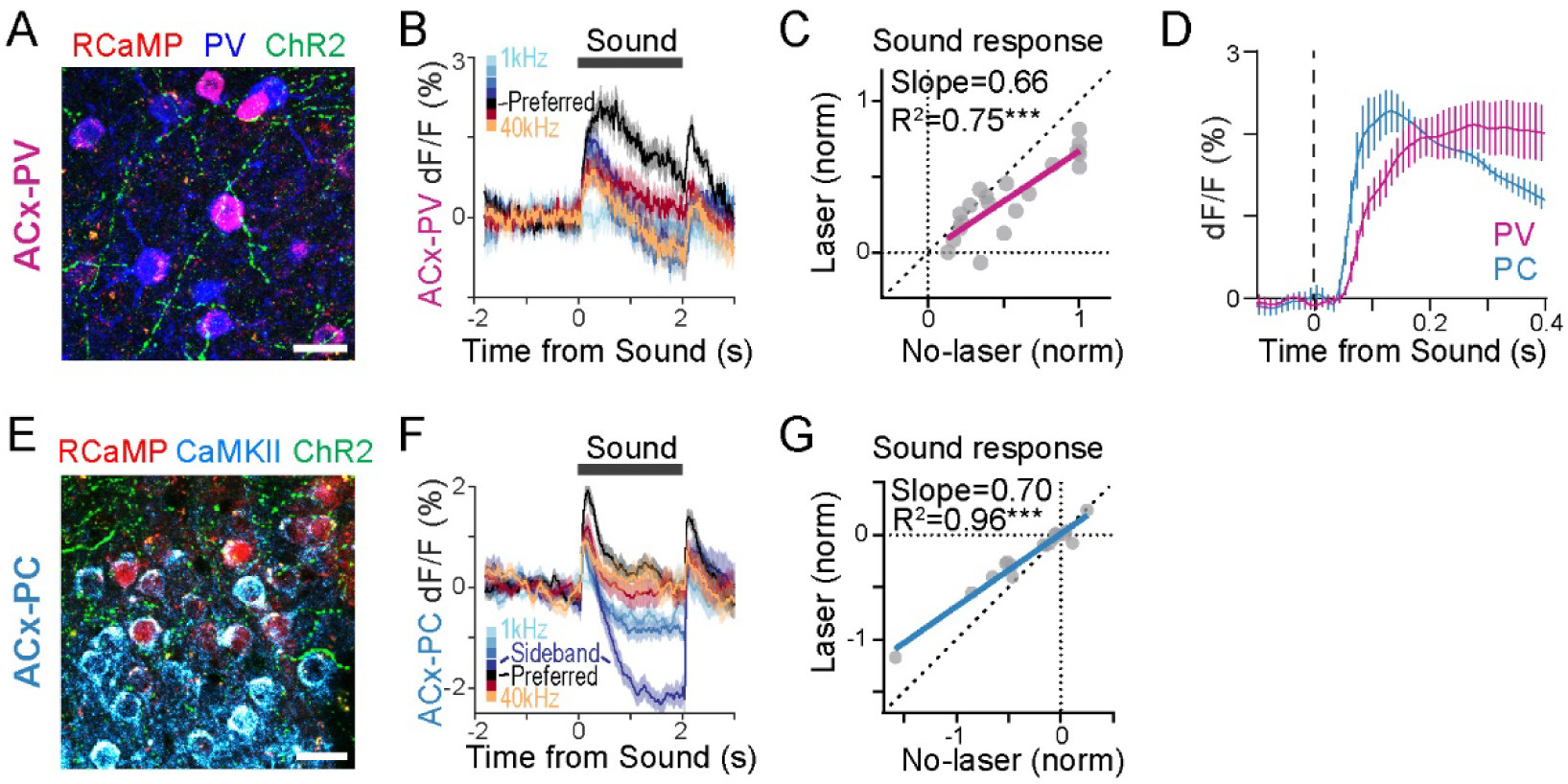
Activation of BF-PV neurons inhibits auditory responses of PV inhibitory neurons while disinhibits principal neurons, related to Figure 7. (A) RCaMP-expressing (red) PV neurons (anti-PV, blue) and ChR2 (EYFP tagged, green)-expressing BF-PV innervations in the auditory cortex. Scale bar, 25μm. (B) Auditory PV neurons’ (ACx-PV) responses to different tone frequencies in an example animal. (C) Optogenetic activation of BF-PV neurons produces greater suppression effects for the tone frequencies with bigger sound responses in ACx-PV neurons. The tone frequencies with no obvious sound response (< 10% of peak response at the preferred frequency) are excluded [R2 = 0.75, p < 0.0001, Pearson]. (D) Comparison of the peak sound responses of auditory PV (ACx-PV) [n = 4 mice] and principal neurons (ACx-PC) [n = 3 mice]. (E) RCaMP-expressing (red) principal neurons (anti-CaMKII, cyan) and ChR2 (EYFP-tagged, green)-expressing BF-PV innervations in the auditory cortex. Scale bar, 25μm. (F) Auditory principal neurons’ (ACx-PC) responses to different tone frequencies in an example animal. Note the post-peak suppression (trough) at sideband frequencies. (G) Optogenetic activation of BF-PV neurons produces greater disinhibition effects for the tone frequencies with deeper troughs in ACx-PC neurons. The tone frequencies with no obvious sound response are excluded [R2 = 0.96, p < 0.0001, Pearson]. Data are represented as means ± SEM. *p < 0.05, **p < 0.01, ***p < 0.001.

**Figure S7.**
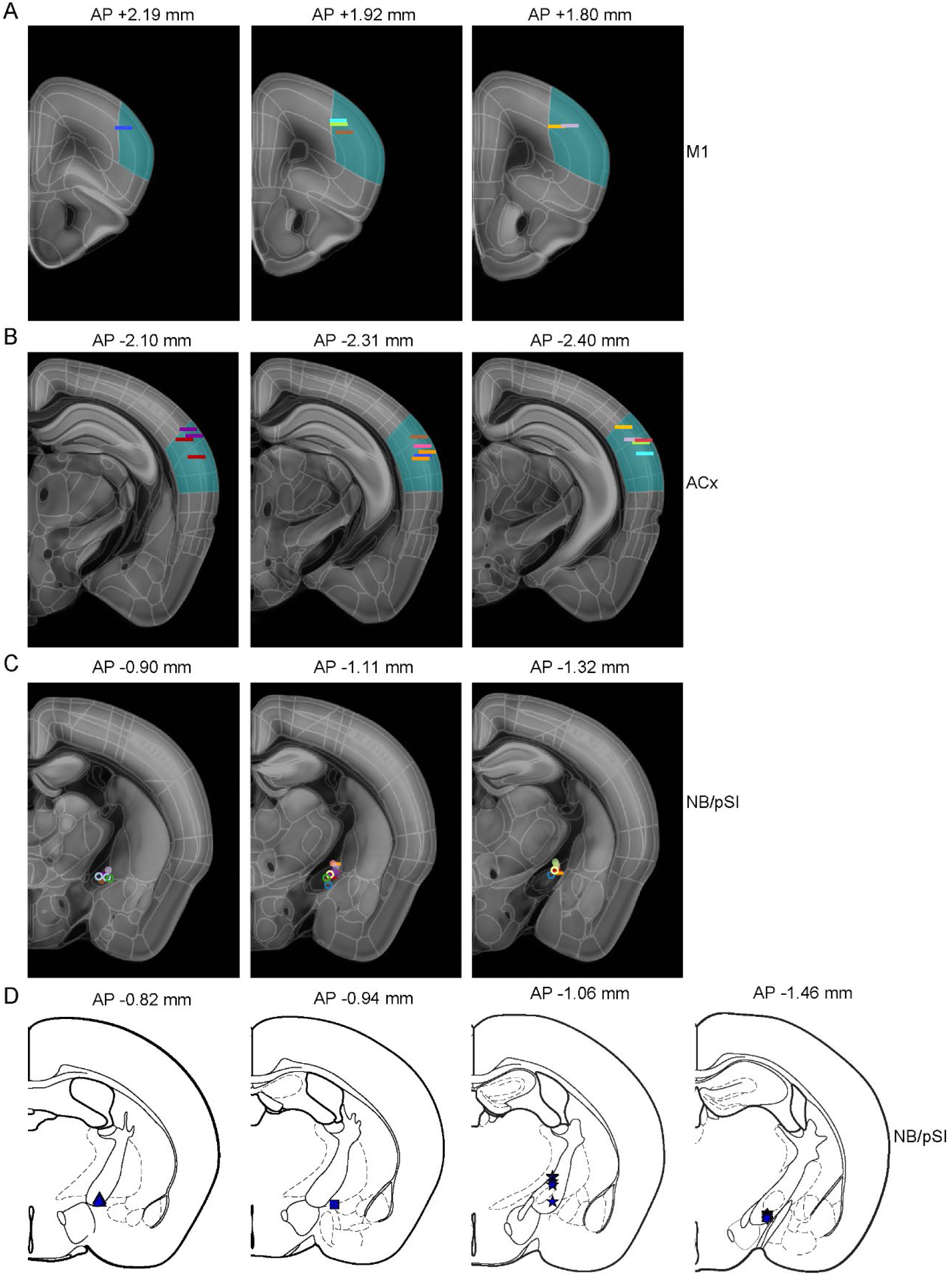
Fiber implantation sites for photometry and optogenetic experiments, related to Figure 2, 3, 4, 6 & 7. (A-B) Schematic summary of the location of optic fibers (400μm) used for photometry recordings in the primary motor cortex (A) and auditory cortex (B) in Figure 2, 3, 4 & 7. Aim regions are highlighted. Each line represents one fiber tip site. Each color represents one animal. Symbols with the same color represent bilateral fiber implantation. (C) Schematic summary of the locations of optic fibers (200μm) used for optogenetic manipulation in NB/pSI regions. Each fiber tip site is represented as a short line for experiments in Figure 7, a solid dot for opto-activation, or an open dot for opto-inhibition in Figure 6. Each color represents one animal. (D) Schematic summary of spike recording positions of identified neurons in Figure 2 & 4. Different symbols represent individual animals.

